# *Tmed2* regulates Smoothened trafficking and Hedgehog signalling

**DOI:** 10.1101/2020.04.20.049957

**Authors:** Giulio Di Minin, Charles E. Dumeau, Alice Grison, Wesley Chan, Asun Monfort, Loydie A. Jerome-Majewska, Anton Wutz

## Abstract

Hedgehog (HH) signalling plays a key role in embryonic pattering and stem cell differentiation. Compounds that selectively bind Smoothened (SMO) can induce cell death in mouse embryonic stem cells (ESCs). Here we perform a genetic screen in haploid ESCs and discover that SMO inhibits a cell death pathway that resembles dissociation induced death of human ESCs and Anoikis. In mouse ESCs, SMO acts through a G-protein coupled mechanism that is independent of GLI activation. Our screen also identifies the Golgi proteins Tmed2 and Tmed10. We show that TMED2 binds SMO and controls its abundance at the plasma membrane. In neural differentiation and neural tube pattering Tmed2 acts as a repressor of HH signalling strength. We demonstrate that the interaction between SMO and TMED2 is regulated by HH signalling suggesting SMO release form the ER-Golgi is critical for controlling G-protein and GLI mediated functions of mammalian HH signalling.

## Introduction

Hedgehog (HH) signalling is crucial for embryonic development and regulates tissue repair (Peng, Frank et al., 2015, Petrov, Wierbowski et al., 2017, Petrova & Joyner, 2014). Consequently, genetic or pharmacologic alterations of HH pathway cause developmental abnormalities. Deregulated HH signalling is also involved in tumour initiation and dissemination in basal cell carcinoma (BCC), medulloblastoma and rhabdomyosarcoma (Pak & Segal, 2016). Compounds targeting the signal transducer Smoothened (*Smo*) are in clinical trials of cancer therapies (Girardi, Barrichello et al., 2019).

Binding of HH ligands releases SMO from the inhibition by the HH receptor Patched-1 (*Ptch1*) (Briscoe & Therond, 2013). SMO then mediates transcription-dependent and independent effects (Kong, Siebold et al., 2019). SMO is a G-protein coupled receptor (GPCR) and regulates a number of cellular processes including the actin cytoskeleton of multiple cell types promoting their motility and migration (Bijlsma, Damhofer et al., 2012, Polizio, Chinchilla et al., 2011a, Yam, Langlois et al., 2009), and calcium levels rewiring cell metabolism in muscles and brown fat (Teperino, Amann et al., 2012). An important function of SMO is the activation of GLI family transcription factors. Activation of GLI proteins in vertebrates is restricted in a specialized plasma membrane compartment, the primary cilium (Corbit, Aanstad et al., 2005). In mammals, SMO translocation to the primary cilium upon HH stimulation is necessary for regulation of GLI2 and GLI3 processing (Milenkovic, Scott et al., 2009, Rohatgi, Milenkovic et al., 2007, Wang, Zhou et al., 2009). In the absence of HH signalling partial proteolysis by the proteasome at the base of the primary cilium generates truncated GLI proteins that enter the nucleus and act as repressors. SMO activation and translocation to the cilia frees GLI proteins from their binding proteins in the tip of the cilia and prevents modifications that are required for the proteolytic cleavage. Intact GLI2 and GLI3 then function as activators of transcription. One of their targets is *Gli1* that further contributes to GLI mediated transcription (Niewiadomski, Niedziolka et al., 2019). The importance of both GLI activators and repressors is observed in neural tube and limb development (Briscoe & Therond, 2013). The central position of SMO in the HH signalling pathway has spurred studies into the mechanisms by which SMO is regulated and coupled to downstream components. This has led to the identification of cross-talk between lipid metabolism and HH signalling. Oxysterols and cholesterol can bind SMO and induce downstream events (Byrne, Sircar et al., 2016, Cooper, Wassif et al., 2003, Huang, Nedelcu et al., 2016, Luchetti, Sircar et al., 2016, Xiao, Tang et al., 2017). PTCH1 has homology to the cholesterol transporter Niemann-Pick C1 and possesses multiple cholesterol binding sites (Gong, Qian et al., 2018, Kowatsch, Woolley et al., 2019, Qi, Di Minin et al., 2019, Qi, Schmiege et al., 2018a, Qi, Schmiege et al., 2018b, Qian, Cao et al., 2019). It has been proposed that PTCH1 regulates SMO activity by preventing its accumulation at the primary cilium (Byrne et al., 2016, Kinnebrew, Iverson et al., 2019, Kong et al., 2019, Luchetti et al., 2016). Compartmentalization facilitates a focused change of a regulatory molecule that can then be sensed to trigger activation of the signalling cascade. SMO localized at the plasma membrane or in endocytic vesicles is prevented to accumulate at the primary cilium by PTCH1 activity, which reduces accessibility of cholesterol or specific derivatives in the ciliary compartment. HH binding to PTCH1 is thought to inhibit its transport activity and leads to increased cholesterol accessibility and lateral transport of SMO into the primary cilia, where SMO can regulate GLI processing (Kinnebrew et al., 2019, Kong et al., 2019, Milenkovic et al., 2009). In contrast, G protein regulation by SMO is not restricted to the primary cilium (Fan, Chen et al., 2014, Myers, Neahring et al., 2017, Pandit & Ogden, 2017, Yuan, Cao et al., 2016). Therefore, the GPCR and GLI processing functions of SMO can be regulated independently in different membrane compartments.

Previous studies have investigated SMO function applying GLI transcriptional reporters and genetic screens in 3T3 fibroblasts using siRNA (Jacob, Wu et al., 2011) and CRISPR-Cas9 (Breslow, Hoogendoorn et al., 2018, Kinnebrew et al., 2019, Pusapati, Kong et al., 2018) libraries. These studies have provided a detailed understanding of pathways for the assembly of and trafficking to the primary cilia, and identified factors for processing of GLI proteins. However, the molecular mechanism of SMO regulation outside the ciliary compartment remains less well understood. SMO is translated and imported into the ER membrane (Denef, Neubuser et al., 2000, Incardona, Gruenberg et al., 2002). Export from the ER, transport through the Golgi, and trafficking to the plasma membrane are highly regulated processes (Dong, Filipeanu et al., 2007). The early secretory pathway involves bidirectional transport between the ER and the Golgi compartments that is mediated by coat protein complex I (COPI)-coated and COPII-coated vesicles (Brandizzi & Barlowe, 2013). Retrograde transport has been implicated in quality control of membrane proteins before they reach the plasma membrane. Following ER exit proteins are further modified in the Golgi, and subsequently sorted to different membrane compartments of the cell. How SMO shuttles from the ER to the plasma membrane remains unknown. The abundance at the plasma membrane is critical for the signalling strength of GPCRs (Dong et al., 2007).

## Results

### Screening for resistance to SMO selective compounds in haploid ESCs identifies Anoikis modulators

When investigating the effect of the HH signalling on mouse ESCs, we observed that Smoothened agonist (SAG) and Purmorphamine (PMP) decreased survival (Fig.1A and Supplementary Fig.1A). In contrast, the HH antagonists KAAD-cyclopamine and SANT1 had no effect on ESCs at concentrations that inhibit GLI transcriptional activity in NIH-3T3 cells (Supplementary Fig.1B). Similarly, recombinant SHH did not affect ESC viability (Fig.1A) at concentrations that induced a strong GLI transcriptional response in NIH-3T3 cells (Supplementary Fig.1B). Furthermore, SAG and PMP were not toxic to NIH-3T3 cells (Supplementary Fig.1C).

High concentrations of SAG (2,5-5 μM) and PMP (5-10 μM) were necessary to promote ESCs death (Fig. 1A) even if their EC50 for HH pathway activation is several orders of magnitude lower at 1-5 nM and 0.1-0.5 μM, respectively (Chen, Taipale et al., 2002, Frank-Kamenetsky, Zhang et al., 2002, Wu, Walker et al., 2004).

**Figure 1.**
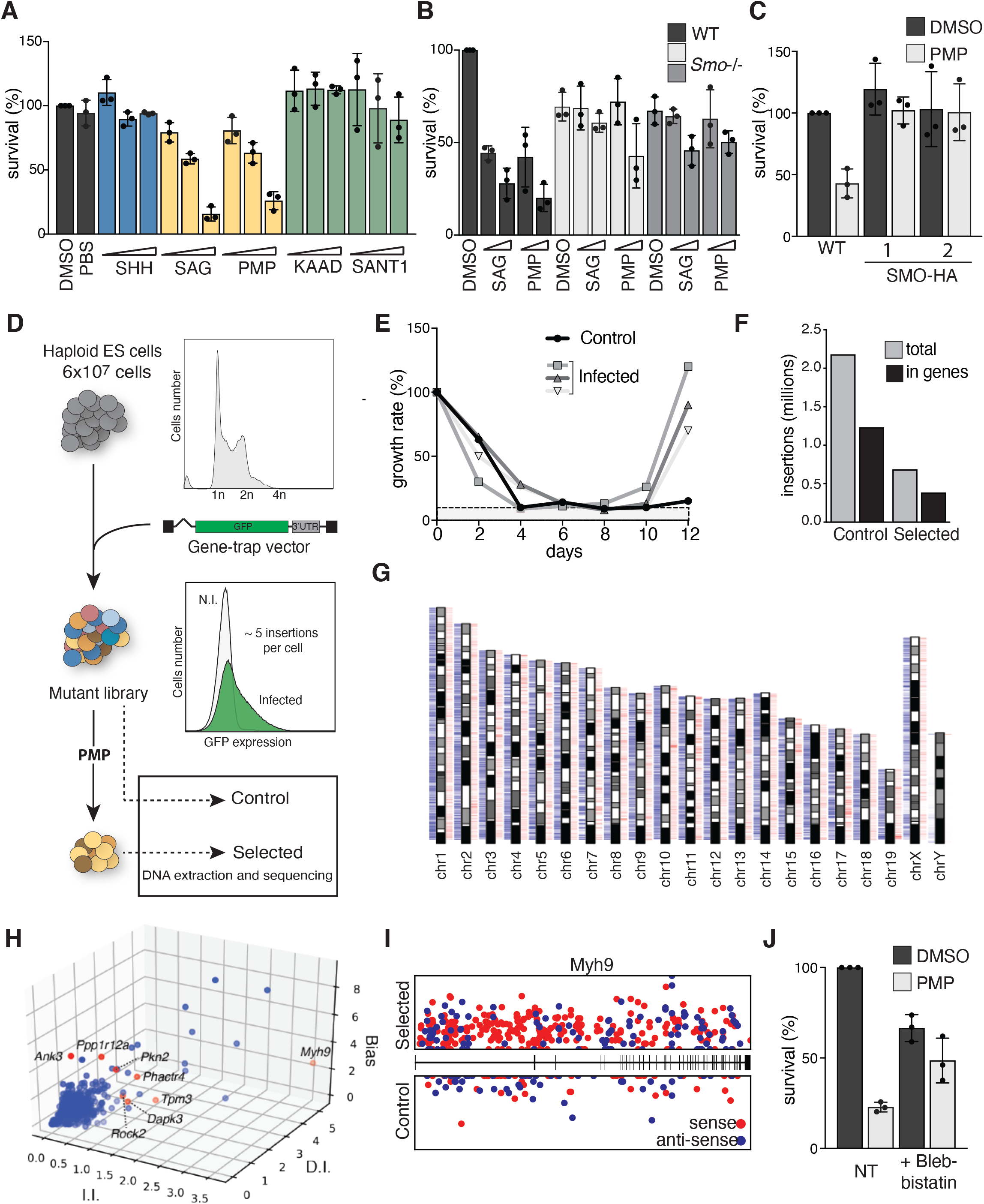
Genetic screen in haploid ESCs for PMP resistance identifies Anoikis modulators. **(A)** SAG and PMP decrease survival of ESCs treated for 48h with SHH (50-500 ng/ml), SAG (0.5-5 μM), PMP (1-10 μM), Cyclopamine-KAAD (1-5 μM), and SANT-1 (10-50 μM). Cell counts are normalized to DMSO treated sample. **(B)** SAG and PMP cytotoxic effects are reduced in *Smo*−/− ESCs. Survival of *Smo*−/− and control ESCs treated for 48h with SAG (2.5-5 μM) and PMP (5-10 μM). The effect on the two independent clones described in Supplementary Fig. 1D is shown and compared to control cells. **(C)**SMO-HA overexpression confers resistance to PMP. Survival of ESCs after PMP treatment (5 μM). The effect on two independent clones described in Supplementary Fig. 1E is shown. **(D)** Schematic screening strategy for PMP resistance. **(E)** Selection of ESCs resistant to PMP. Infected (three independent experiments) and control cells were treated for 12 days with PMP. ESCs were passed and counted every 2 days. The graph shows the percentage of the counted cells relative to the number of cells plated on day 0. The dashed line indicates a baseline of irradiated MEFs used for ESC culture. **(F)** Depiction of insertion numbers genome wide (grey) and in gene transcription units (black), for control and PMP selected samples. **(G)** Chromosomal distribution of insertions in control (blue, above) and selected (red, below) samples are shown. **(H)** Identification of genes conferring PMP resistance by enrichment of independent viral insertions (I.I.), disrupting insertions (D.I.) and gene trap orientation bias (Bias). Top candidates implicated in anchorage independent growth are marked in red and annotated. **(I)** Distribution of I.I. within *Myh9* in PMP selected (top) and control (bottom) samples. Gene trap insertions in the orientation of the gene transcription unit (sense) are marked in red, and antisense insertions are marked in blue. **(J)** The MYH9 inhibitor Blebbistatin mediates resistance to PMP. Survival of ESCs treated for 48h with PMP with or without the addition of Blebbistatin.

To confirm the SMO dependency on this phenotype, we derived *Smo−/−* ESCs using CRISPR/Cas9 nucleases (Supplementary Fig. 1D) and tested their response to this class of compounds. *Smo* mutant cells were considerably more resistant to SAG and PMP treatment compared to control cells (Fig. 1B). In addition, we observed that the growth rate of *Smo−/−* ESCs was decreased in untreated conditions compared to control cells (Fig. 1B). This phenotype was particularly pronounced in a chemically defined culture medium that maintains ground state pluripotency of ESCs (Ying, Wray et al., 2008).

We next investigated if SAG and PMP toxicity could be mediated by hyperactivation of SMO and HH signalling. For this we stably over-expressed a SMO-HA construct in mouse ESCs (Supplementary Fig. 1E). Notably, SMO-HA over-expression had little effect on ESC cultures even though high levels of SMO-HA were detected by Western analysis (Fig. 1C). Instead, increased SMO expression had a protective role from PMP cytotoxicity (Fig. 1C). Taken together these observations show that the effect of SAG and PMP on ESC survival is dependent on SMO. However, the observed protective effect of SMO also raises the question by which mechanism SMO functions in ESCs.

To further characterize the effect of SAG and PMP, we performed a genetic screen in haploid ESCs. Haploid ESCs have been previously shown to be suitable for efficient screening of developmental pathways (Leeb, Dietmann et al., 2014, Monfort, Di Minin et al., 2015, Yilmaz, Peretz et al., 2018) and the sensitivity of ESCs to PMP provided a selection strategy. We infected 60 million haploid ESCs with a viral gene trap vector to obtain a genome-wide library of mutations (Fig. 1D). Three independent mutant pools were subsequently split into two populations that were either selected with PMP or used as a control. We observed a PMP resistant population after 12 days (Fig. 1E). Over 2 million independent viral insertions were identified by next generation sequencing (NGS) in control and selected samples. Insertions were distributed over all chromosomal regions and showed an expected bias in transcribed regions (Fig. 1F-G). Candidate gene prediction was performed using the HaSAPPy package (Di Minin, Postlmayr et al., 2018) (Fig. 1H and Supplementary Fig. 1F). We did not detect enrichment of known genes associated with the canonical HH pathway or primary cilium in our screen (Supplementary Table1). Instead genes involved in cytoskeleton dynamics and anchorage independent growth were strongly enriched (Fig. 1H and Supplementary Table1). Mutations in these genes are frequently identified in tumours conferring resistance to Anoikis-induced cell death (Supplementary Fig. 1G). We detected strong evidence for selection of inactivating mutations of *Myh9* in PMP selected ESCs (Supplementary Table1 and Fig. 1I) and the specific MYH9 inhibitor Blebbistatin rescued mouse ESCs from PMP induced death (Fig. 1J). This observation indicates that MYH9 activity is required for cytotoxicity of SMO selective compounds in ESCs.

### SMO counteracts dissociation-induced apoptosis in ESCs

Hyperactivation of MYH9 is the direct cause of dissociation-induced apoptosis in human ESCs (Chen, Hou et al., 2010, Ohgushi, Matsumura et al., 2010). After passaging, control ESCs attached and spread on the cell culture plate (Fig. 2A). In contrast, PMP treated cells became motile, displayed blebbing of the plasma membrane (Fig. 2B), and ultimately formed apoptotic bodies. Annexin V staining confirmed that SAG and PMP induced apoptosis in ESCs (Fig. 2C). Apoptosis upon dissociation of human ESCs can be suppressed by inhibitors of ROCK kinase (Chen et al., 2010, Ohgushi et al., 2010). We find that ROCK inhibition also protected mouse ESCs from PMP induced apoptosis (Fig. 2D-E).

**Figure 2.**
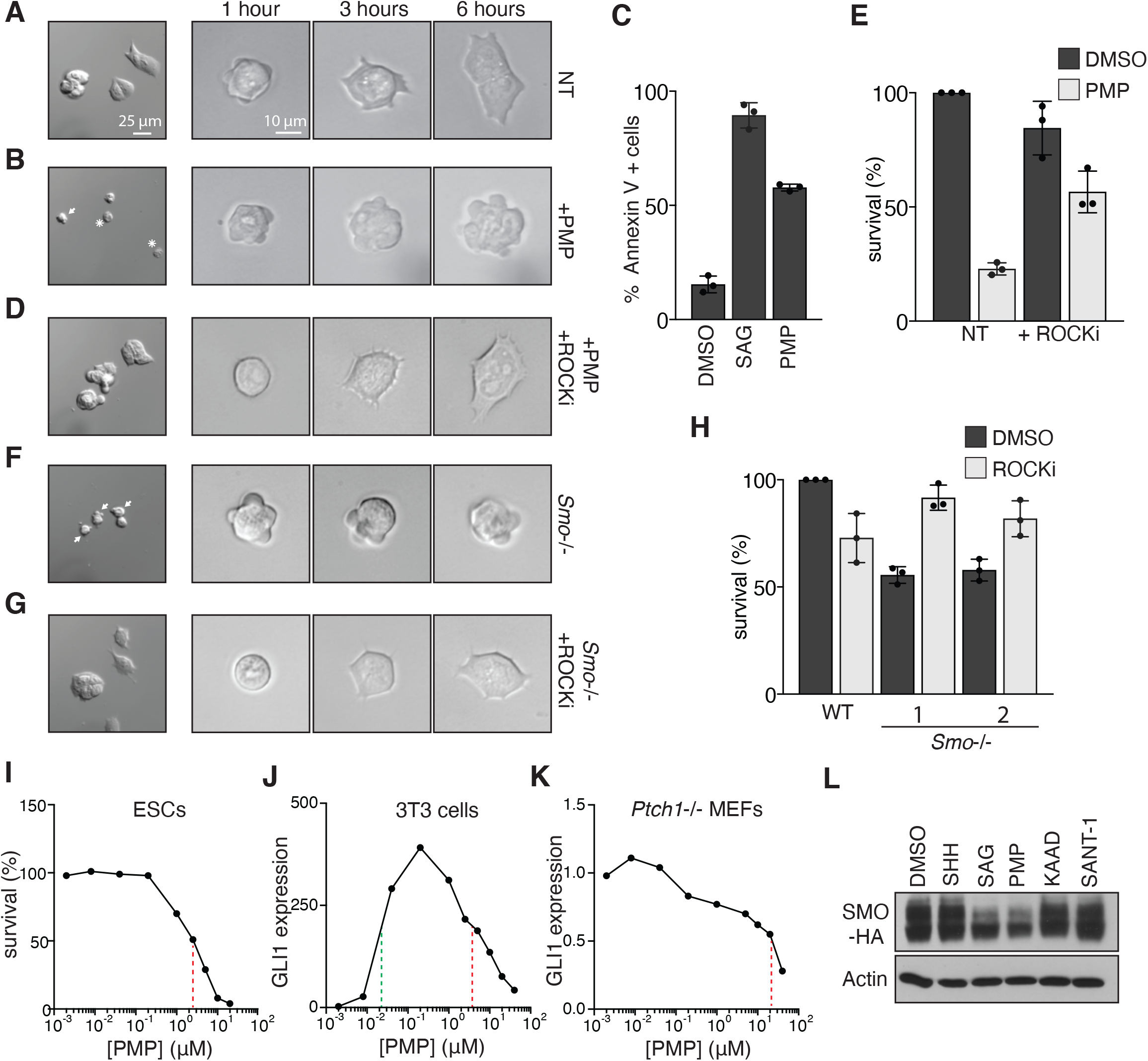
SMO supports survival of ESCs after dissociation. **(A-B)** PMP induces blebbing and death after dissociation of ESCs. On the left, ESCs morphology 24h after dissociation and plating on Matrigel (A, untreated; B, pre-treated with PMP for 24h). Cells showing membrane blebbing (arrow), and apoptotic bodies (asterisk) are indicated. Images at 1, 3, and 6 hours after plating are shown on the right. **(C)** Apoptosis after SAG and PMP treatment measured by Annexin V staining. **(D-E)** ROCK inhibition prevents PMP induced apoptosis. **(D)** Images of ESCs treated with PMP and ROCK inhibitor Y-27632 (ROCKi) as in A. **(E)** Survival of ESCs treated for 48h with PMP with or without the addition of ROCKi. **(F-H)** *Smo* mutation affects ESC survival after dissociation. Images of *Smo*−/− ESCs as in A untreated **(F),** and treated with ROCKi **(G)**. **(H)** Survival of *Smo*−/− ESCs after 48h with or without addition of ROCKi. **(I)** Dissociation induced apoptosis is prompted by high PMP concentrations. Survival of ESCs treated for 48h with different PMP concentrations (2 nM - 20 μM). Red dotted line highlight IC50 at 2.5 μM **(J-K)** High concentrations of PMP repress GLI activity and HH signalling. **(J)** RT-qPCR of *Gli1* mRNA in NIH-3T3 cells treated with increasing concentrations of PMP (2 nM - 40 μM) and normalized to untreated samples. Dotted lines mark PMP EC50 (green, at 30 nM) and IC50 (red, at 4 μM). **(K)** RT-qPCR of *Gli1* mRNA in *Ptch1*−/− MEFs treated with increasing concentrations of PMP (2 nM - 40 μM) and normalized to untreated samples. Red dotted line highlight IC50. **(L)** Effects of HH pathway targeting compounds on SMO stability. ESCs expressing SMO-HA were treated for 24h with the indicated compounds and SMO levels were analysed by immunoblot using an HA antibody. Actin is shown as loading control.

Similar to PMP treated ESCs, *Smo* mutant ESCs showed pronounced membrane blebbing (Fig. 2F). Thereby decreased survival after dissociation could also explain the lower growth rate of *Smo* mutant ESCs (Fig. 1B). Conversely, ROCK inhibition promoted *Smo* mutant ESCs survival (Fig. 2G-H). Taken together these data show that SAG and PMP affect ESC survival similar to a *Smo* mutation.

Cytotoxicity in ESCs required a PMP concentration above 2.5 μM (Fig. 2I) that is an order of magnitude higher than the concentration used to activate SMO. We verified that in NIH-3T3 cells high PMP concentrations repress GLI transcription with an IC50 approximately of 4 μM (Fig. 2J), which is consistent with previous studies. Concentrations above 1μM become inhibitory rather then activating leading to a biphasic bell-shaped activity curve (Chen et al., 2002, Frank-Kamenetsky et al., 2002, Wu et al., 2004). We further confirmed these results in *Ptch1*−/− MEFs where absence of PTCH1 leads to constitutive activation of the HH pathway (Fig. 2K). Addition of PMP at concentrations higher than 2.5 μM strongly repressed GLI transcription in *Ptch1*−/− MEFs.

Our observations in ESCs could be explained by considering that high concentrations of SMO agonists acted inhibitory and phenocopied a SMO mutation. The evidence that increasing SMO levels alleviated PMP toxicity (Fig. 1C) suggested that a decrease in SMO protein stability could contribute to cell death. In support of this notion, we find that SAG and PMP treatment reduced SMO protein in ESCs (Fig. 2L). Taken together our results indicate that *Smo* functions to prevent death in pluripotent cells when cell-cell contact is interrupted. Although a previous study did not observe general toxicity of high SAG concentrations (Chen et al., 2002), we cannot fully rule out additional effects independent of SMO. However, we observe similar effects with the structurally unrelated SMO selective compounds SAG and PMP.

### SMO supports ESC survival through modulating Rac1/RhoA activity

SAG and PMP treatment reduced survival but did not change *Gli1* expression in ESCs suggesting an GLI-independent mechanism (Fig. 3A). It has been shown that SMO regulates heterotrimeric G proteins of the G_i_ subclass (Brennan, Chen et al., 2012, Ogden, Fei et al., 2008, Riobo, Saucy et al., 2006, Shen, Cheng et al., 2013). Direct inhibition of G_i_ proteins by Pertussis toxin (PTX) decreased survival (Fig. 3B) and affected normal spreading of ESCs after dissociation (Fig. 3C). The effects of PTX could furthermore be rescued by ROCK inhibition (Fig. 3B-C) showing that G_i_ protein inhibition induced cell-death similar to PMP. Therefore, SMO could be required in ESCs for G_i_ activation. A known target of SMO inhibition is adenylyl cyclase and PKA activity (Polizio et al., 2011a, Polizio, Chinchilla et al., 2011b). However, PKA activation by Forskolin treatment had no effect on ESC survival (Supplementary Fig. 2A) as reported before (Takahashi, Lee et al., 2014).

**Figure 3.**
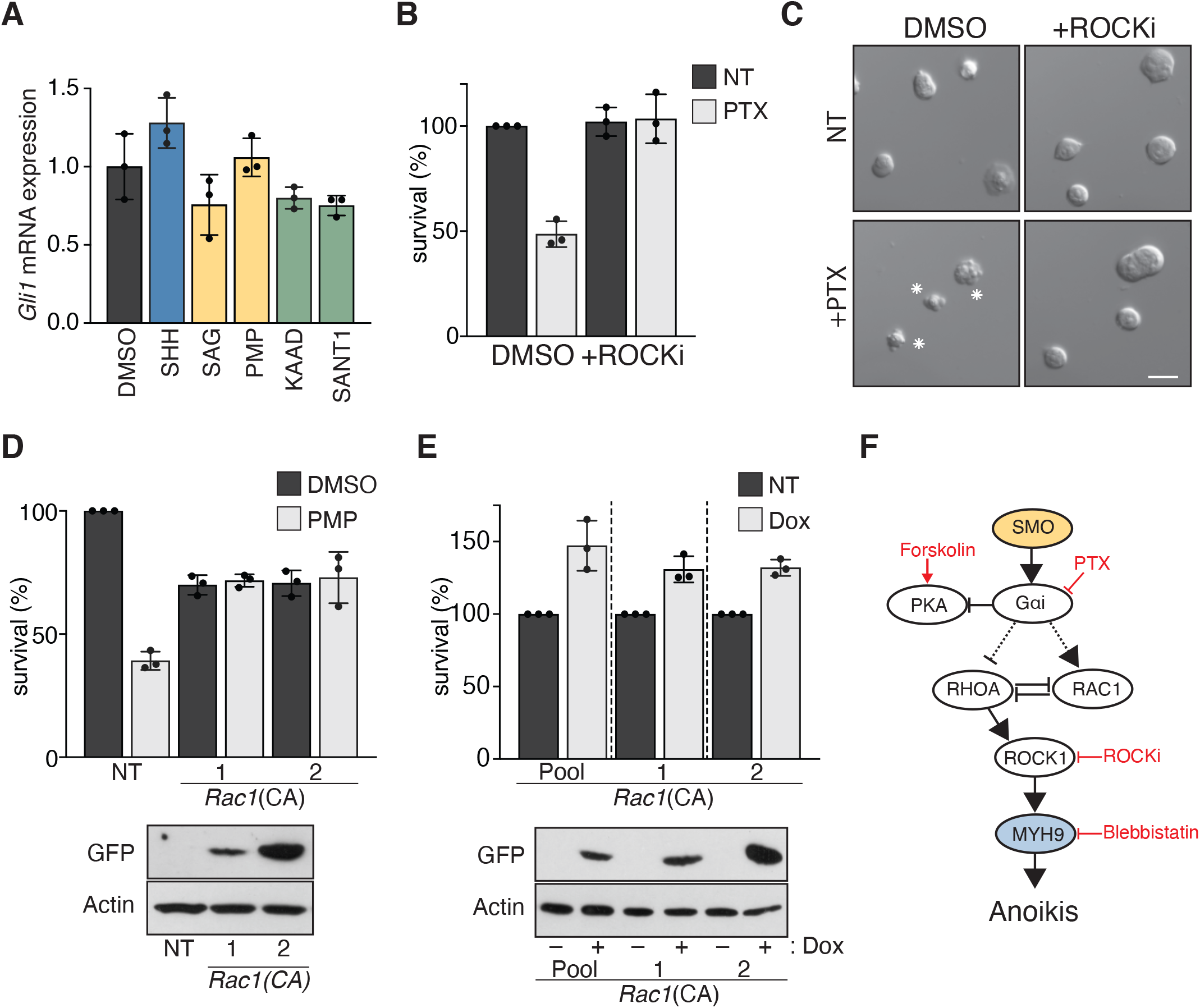
SMO sustains survival of ESCs through activation of G_i_ proteins. **(A)** GLI transcriptional remains unchanged in ESCs treated with HH pathway targeting compounds as in Fig. 1A. ESCs were treated for 24h as indicated, and *Gli1* mRNA levels were measured by qPCR. **(B-C)** G_i_ protein inhibition induces apoptosis in ESCs. Survival of ESCs after 48h treatment with the G_i_ specific inhibitor Pertussis toxin (PTX) with or without the addition of ROCKi. Cell counts are normalized to DMSO treated samples. **(C)** ESC morphology 24h after dissociation. Cells were pre-treated with PTX and ROCKi. Apoptotic bodies are marked by an asterisk. Bars represent 20 μm. **(D)** Constitutive active *Rac1*(CA) confers PMP resistance. Two independent ESCs lines expressing the Rac1(CA)-IRES-GFP cassette were treated with PMP for 48h. Cell survival was normalized to DMSO treated WT ESCs. Immunoblot analysis (below) showing expression of Rac1(CA)-IRES-GFP. **(E)** *Rac1*(CA) increases survival after passaging SMO−/− ESCs. A Rac1(CA)-IRES-GFP construct under the control of a Doxycycline (Dox) inducible promoter was integrated in ESCs. The survival of the transfected cell pool and of two independent clones was analysed with and without Dox induction for 48h. Cell survival was normalized to uninduced cells (NT). Immunoblot analysis (below) showing the induction of the Rac1(CA)-IRES-GFP construct after Dox treatment. **(F)** Model summarizing the mechanism of SMO function in sustaining ESCs survival. Compounds are annotated in red.

In our screen we also identified *Trim71* and *Csde1* that were previously implicated in RAC1 regulation (Supplementary Fig. 2B-D)(Dai, Ma et al., 2018, Wurth, Papasaikas et al., 2016, Xiong, Zhao et al., 2017). Expression of constitutive active RAC1(Q61L) increased resistance of ESCs to PMP (Fig. 3D). Furthermore, inducible expression of RAC(Q61L) significantly increased the survival of *Smo* mutant ESCs (Fig. 3E). Taken together our data suggest that SMO increases RAC1 activity, which in turn counteracts ROCK1 induced hyper-phosphorylation of myosin and contractility (Fig. 3F). *Smo* mutant ESCs show attachment defects and reduced survival after passaging. Similar defects are observed after treatment with inactivating PMP concentrations and can be rescued by overexpression of SMO. Classic SMO antagonists including KAAD-Cyclopamine and SANT-1 did not affect SMO functions in ESCs. Although, inhibition of SMO GPCR activity has been reported (Polizio et al., 2011a), other studies found that cyclopamine can promote G protein activation by SMO (Teperino et al., 2012). While SMO antagonist specifically act in repressing GLI transcription, their effect on SMO GPCR function is currently less clear. Our observations reveal a new function of SMO for sustaining mouse ESCs survival independent of GLI regulation.

### The Golgi protein TMED2 modulates HH signalling

Our screen identified a second group of candidates with strong enrichment for proteins that are localized in the Golgi, and function in vesicle trafficking (Fig. 4A, Supplementary Fig. 3A and Supplementary Table 1). Among these candidates the p24 family receptors *Tmed2* and *Tmed10* showed the strongest evidence for selection (Fig. 4B). Mutations in other members of the p24 family (Pastor-Cantizano, Montesinos et al., 2016) were not enriched in our screen (Supplementary Fig. 3B). We generated *Tmed2* and *Tmed10* mutations in ESCs using the CRISPR/Cas9 nucleases (Supplementary Fig. 3C and Fig. 4C). Western analysis confirmed the absence of protein in the respective mutant cell lines (Fig. 4C). In addition, the *Tmed10* mutation also strongly reduced TMED2 protein likely due to interdependence of p24 family proteins as observed before (Hou, Gupta et al., 2017, Hou & Jerome-Majewska, 2018, Vetrivel, Gong et al., 2007). Loss of *Tmed2* and *Tmed10* did not measurably affect self-renewal of ESCs (Supplementary Fig. 3D-E). Furthermore, *Tmed2* mutant ESCs showed overall normal Golgi staining (Fig. 4D). These observations are consistent with studies in other cell types (Hou et al., 2017, Hou & Jerome-Majewska, 2018). As expected, *Tmed2* and *Tmed10* mutant ESCs were highly resistant to PMP when compared to control ESCs (Fig. 4E). To confirm a potential role of *Tmed2* in HH signalling, we measured GLI transcriptional activity in NIH-3T3 cells. Depletion of *Tmed2* by siRNA transfection as a secondary assay (Supplementary Fig. 3F) significantly increased expression of *Gli1* (Fig. 4F) and *Ptch1* (Supplementary Fig. 3G) in response to SHH ligand. Therefore, *Tmed2* modulates canonical HH signalling in 3T3 cells. Depletion of *Tmed2* disrupted neither the structure of the primary cilia (Supplementary Fig. 3H) nor the recruitment of SMO to the cilium (Fig. 4G and Supplementary Fig. 3I).

**Figure 4.**
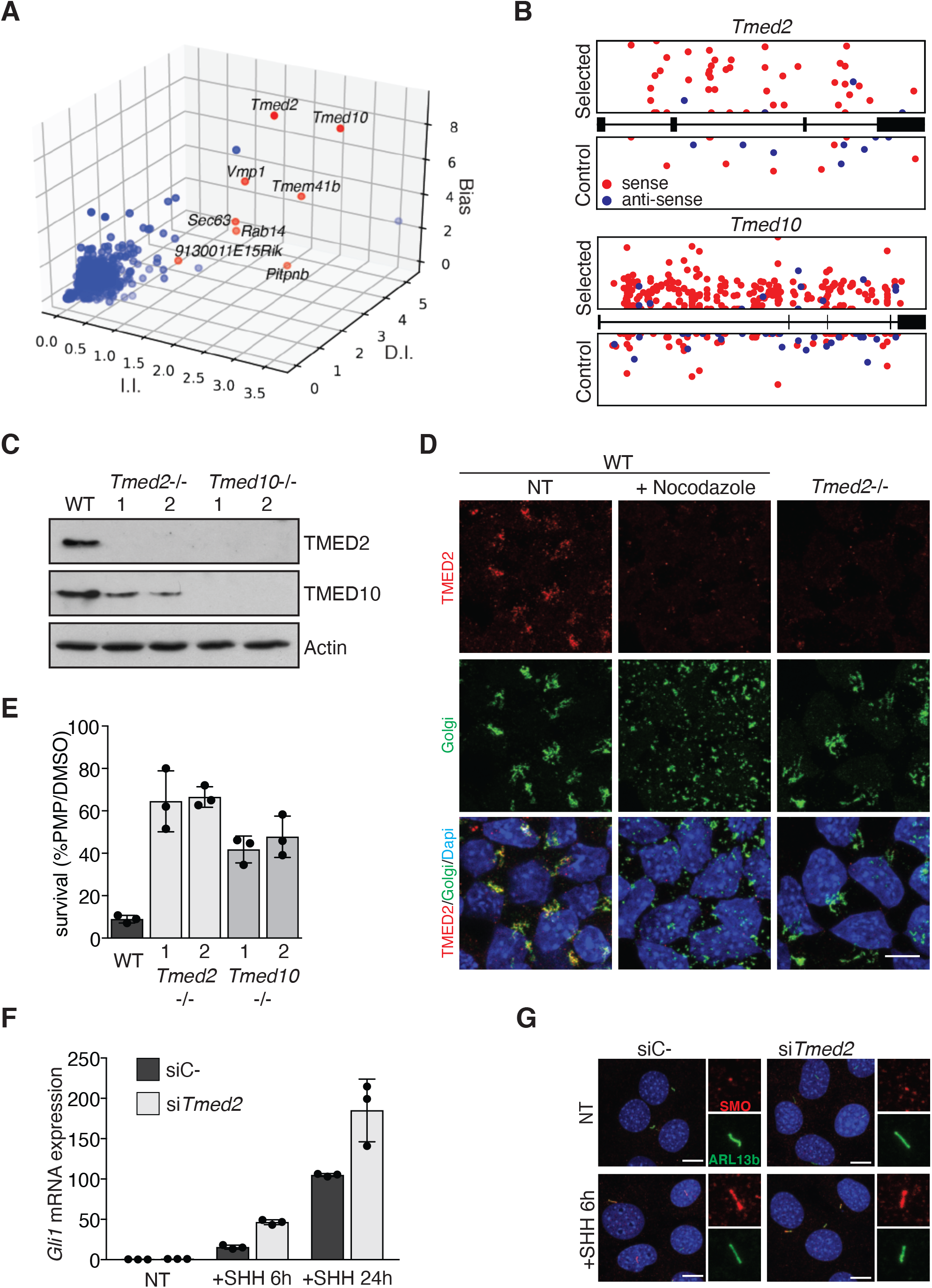
Identification of *Tmed2* as a new modulator of HH signalling. **(A)** Candidates from the PMP resistance screen with reported ER-Golgi localization. Graph shows enrichment of independent viral insertions (I.I.), disrupting insertions (D.I.), and gene trap orientation bias (Bias). Top candidates with reported ER-Golgi localization are marked in red and annotated. **(B)** Distribution of I.I. within *Tmed2* and *Tmed10* in PMP selected (top) and control (bottom) samples. **(C)** Western analysis of parental control (WT), and two clones of *Tmed2*, and *Tmed10* mutant ESCs. **(D)** Immunofluorescence staining of TMED2 and RCAS1 to visualize Golgi structure in WT and *Tmed2*−/− ESCs. ESCs treated with Nocodazole show a disrupted Golgi apparatus; scale = 10 μm. **(E)** *Tmed2* and *Tmed10* mutations confer PMP resistance. Survival of *Tmed2* and *Tmed10* mutant ESCs after PMP treatment. **(F-G)** *Tmed2* knock-down promotes GLI activity after SHH treatment in NIH-3T3 cells. NIH-3T3 cells were transfected with an siRNA targeting *Tmed2* (si*Tmed2*) or a non-targeting control (siC-), and treated with SHH for 6 and 24 hours. **(F)** RT-qPCR of *Gli1* mRNA. **(G)** Immunofluorescence staining of SMO (red) and ARL13B (green) to visualize primary cilia after 6h of SHH treatment. Scale = 10 μm. Inserts (right) magnify SMO and ARL13B localization at the primary cilium.

### *Tmed2* is a negative regulator of HH signalling in neuronal differentiation

To investigate the role of *Tmed2* in development we analysed dorso-ventral neural tube patterning (Dessaud, McMahon et al., 2008). Immunofluorescence staining of neural tube sections of mouse E9.5 embryos showed TMED2 expression in neural progenitors (NPCs) that overlapped with Golgi staining (Fig. 5A and Supplementary Fig. 4A).

**Figure 5.**
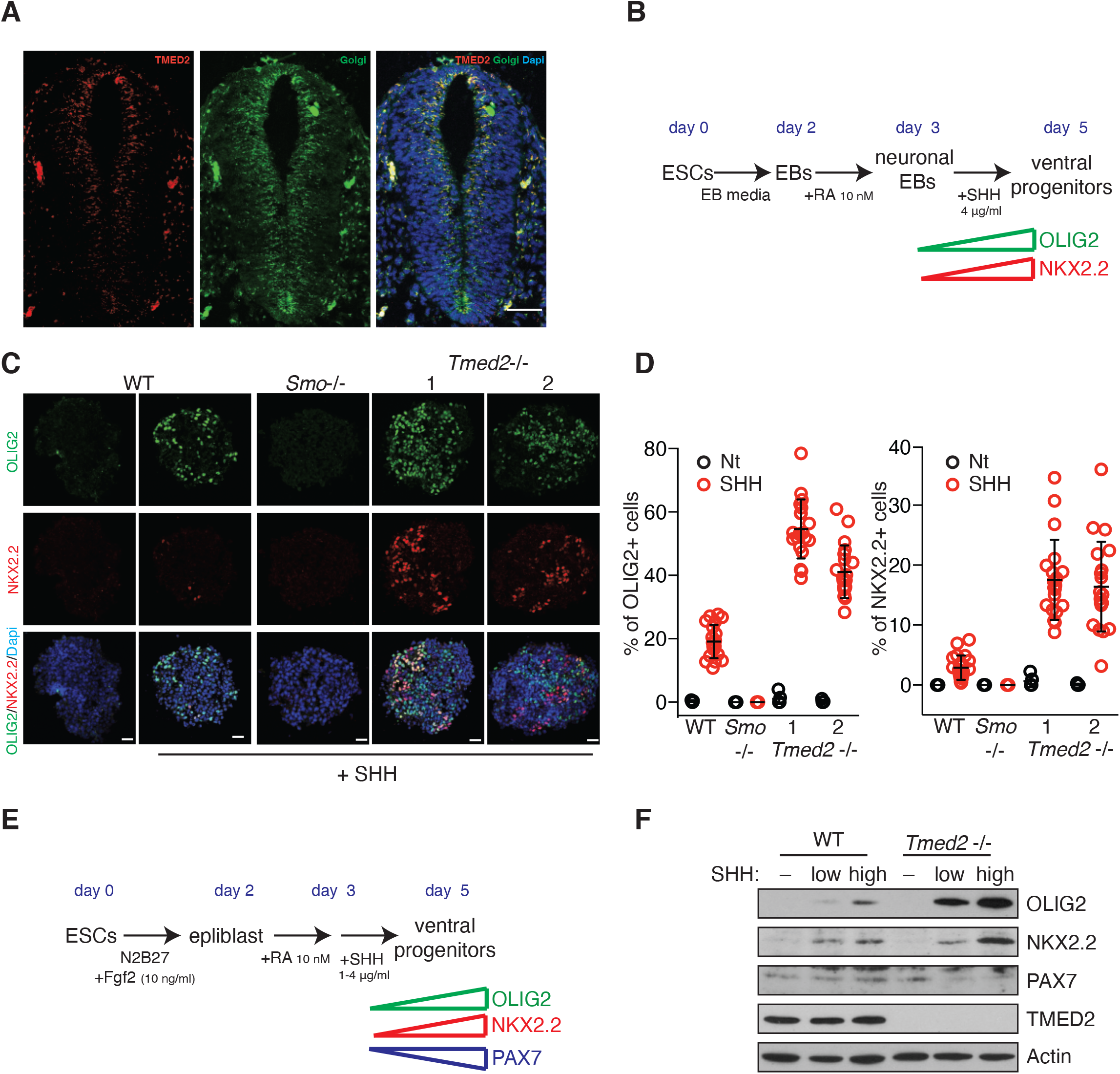
*Tmed2* is a negative regulator of HH signalling in neuronal differentiation. **(A)** Immunostaining of TMED2 (red) and the Golgi marker RCAS1 (green) in neural tube sections of WT E9.5 mouse embryos. Scale = 50 μm. **(B-D)** *Tmed2* mutation increases SHH dependent ventral marker expression in NPCs. **(B)** Schematic overview of the protocol for deriving neuralized EBs from ESCs. The expected timing of NPC markers expression is indicated (bottom). **(C)** Representative immunofluorescence images of ventral markers OLIG2, and NKX2.2 in neuralized EBs of indicated genotypes treated with SHH or not. Scale = 25μm. **(D)** Percentage of cells expressing OLIG2 (left) and NKX2.2 (right) relative to total cell count in neuralized EBs as in C. Individual EBs are plotted (n = 20). **(E-F)** *Tmed2*−/− NPCs derived from ESCs using an adherent culture differentiation (AD) protocol show increased ventral marker expression. **(E)** Schematic overview of the AD protocol indicating the expected timing of NKX2.2, OLIG2, and PAX7 expression (bottom). **(F)** Western analysis of ventral markers OLIG2, and NKX2.2, and the dorsal marker PAX7 in WT and *Tmed2*−/− NPCs on day 5.

This observation suggested that *Tmed2* could influence HH signalling in neural development. The *Tmed2* mutation in mice leads to embryonic lethality before midgestation and *Tmed2−/−* embryos display abnormalities as well as a developmental delay starting from E7.5 (Jerome-Majewska, Achkar et al., 2010). Although *Tmed2−/−* embryos form a neural tube, their developmental delay affects direct comparison to wild type embryos. We observed that NPCs could form normally from *Tmed2* mutant ESCs. Therefore, we decided to initially study *Tmed2* function in dorso-ventral NPCs pattering in culture. We analysed the expression of ventral marker genes OLIG2 and NKX2.2 in NPCs derived by aggregation of ESCs into neuralized embryo bodies (Fig. 5B). The number of *Tmed2−/−* NPCs expressing ventral markers was increased compared to wild type control NPCs in the presence of SHH (Fig. 5C-D). A ventralizing effect was recapitulated when *Tmed2* mutant ESCs were differentiated in NPCs using an adherent culture differentiation (AD) system (Fig. 5E). *Tmed2−/−* NPCs were characterized by increased expression of OLIG2 and NKX2.2 compared to controls after SHH treatment (Fig. 5F). Conversely, the dorsal marker PAX7 was decreased in *Tmed2* mutant NPCs. Our results show that a *Tmed2* mutation enhances HH signalling *in vitro*, and suggest *Tmed2* as a negative regulator of HH signalling in neural differentiation.

### *Tmed2* regulates HH signalling in development

During embryo development, HH expression from the floor plate induces *Nkx6.1* in ventral NPCs. Subsequently, high HH signalling intensity drives the specification of motor neurons that co-express *Olig2*. Neural tubes of E10.5 control embryos appear twice to three times the size of E10.5 *Tmed2−/−* neural tubes (Supplementary Fig. 5A) owing to the well described developmental delay (Fig. 6B). To compensate for this delay and analyse comparable stages of neural tube development, we compared sections of E10.5 *Tmed2−/−* embryos also to E9.0 control embryos. As expected, E9.0 control embryos showed an extensive domain of NKX6.1 expression that contained a confined region of OLIG2 expression (Fig. 6A-C, Supplementary Fig. 5B and Supplementary Table 2). A similar ratio between OLIG2+ and NKX6.1+ progenitors was also observed in E10.5 control embryos (Fig. 6C and Supplementary Fig. 5A). *Tmed2−/−* embryos showed an extended and more dorsal OLIG2 expression domain (Fig. 6A-C and Supplementary Fig. 5C). In contrast, the dorsal marker PAX7, which is repressed by the SHH gradient, was confined to a restricted dorsal region in the neural tube of *Tmed2−/−* embryos, whereas PAX7 covered an extensive region in control embryos (Fig. 6D). Despite the overall developmental delay of *Tmed2* mutant embryos, the ventralization of markers in the neural tube is striking and consistent with the view that ventral HH effects are strengthened in the absence of *Tmed2*.

**Figure 6.**
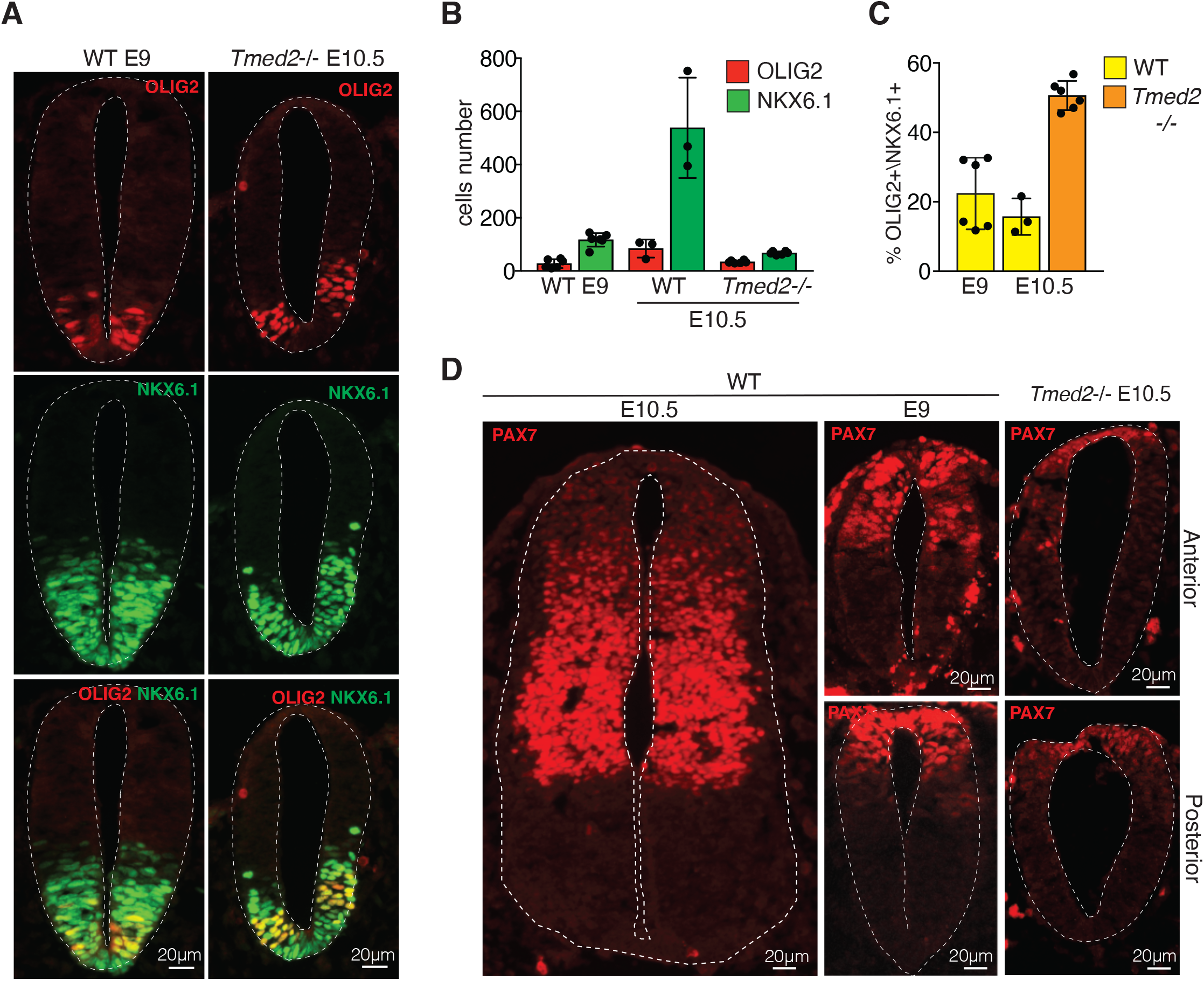
*Tmed2* mutation affects neural tube pattering. **(A-C)** *Tmed2* mutation increases expansion of the OLIG2 expression domain. **(A)** Neural tube sections of E9.0 control and E10.5 *Tmed2*−/− embryos were stained for the ventral markers OLIG2 and NKX6.1. Scale = 20μm. **(B)** Quantification of OLIG2 and NKX6.1 expressing NPCs, and **(C)** the percentage of NPCs expressing OLIG2 relative to NKX6.1 positive NPCs in neural tube section of E9.0 and E10.5 control, and E10.5 *Tmed2*−/− embryos. **(D)** *Tmed2*−/− embryos show decreased PAX7 expression. Neural tube sections derived from E9.0 and E10.5 control, and E10.5 *Tmed2*−/− embryos were stained for the dorsal marker PAX7. Scale = 20 μm. Anterior and posterior sections of the neural tube are shown for E9.0 control, and E10.5 *Tmed2*−/− embryos. The E10.5 WT section (left) is derived from the posterior region of the embryo.

### *Tmed2* restricts SMO abundance at the plasma membrane

The amount of SMO at the plasma membrane is a major determinant of the strength of G protein and GLI mediated effects. *Tmed2* has been implicated in vesicle trafficking and protein secretion (Hou & Jerome-Majewska, 2018, Luo, Wang et al., 2007, Luo, Wang et al., 2011, Stepanchick & Breitwieser, 2010, Sun, Zhang et al., 2018), which suggested a possible role in controlling the amount of SMO that reaches the plasma membrane. To test this hypothesis, we introduced *Tmed2* mutations into ESCs that transgenically express an HA tagged version of SMO (SMO-HA), which can be detected with a specific antibody. Subsequently, we analysed SMO translocation to the plasma membrane in NPCs by using a cell-impermeable biotinylation reagent and purification of biotinylated proteins. A small amount of SMO-HA was detected at the plasma membrane in control cells in the absence of SHH ligand. SHH treatment induced SMO-HA translocation to the plasma membrane (Fig. 7A and Supplementary Fig. 6A). In *Tmed2* mutant cells SMO-HA was abundant at the plasma membrane even without SHH treatment. Importantly, the TMED2 mutation had no effect on the total cellular amount of SMO-HA. Our data therefore demonstrate that TMED2 regulates the abundance of SMO at the plasma. We further noted a lower apparent molecular weight of SMO-HA at the plasma membrane in *Tmed2* mutant cells than in control cells after SHH treatment. This observation indicates that SMO-HA reaches the plasma membrane with different modifications.

**Figure 7.**
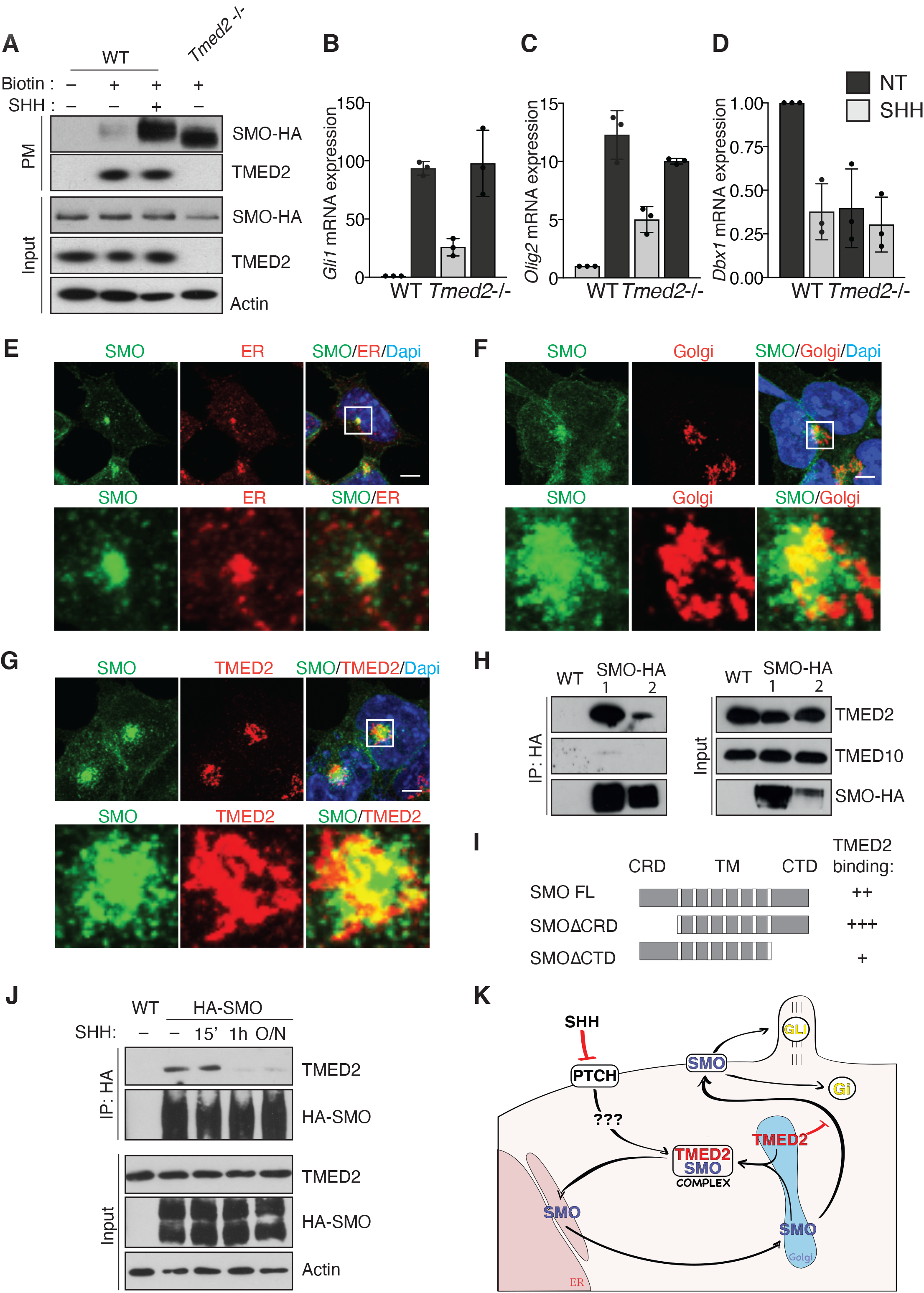
TMED2 binds SMO and regulates its abundance at the plasma membrane in a SHH dependent manner. **(A)** *Tmed2* regulates SMO abundance at the plasma membrane. Cells were treated with NHS-SHHS-Biotin (Biotin) to label plasma membrane proteins, and with SHH as indicated. Western analysis of plasma membrane proteins (PM) and input before purification (1/50 of pull-down) are shown. **(B-D)** qPCR analysis of **(B)** *Gli1*, **(C)** the ventral marker *Olig2*, and **(D) the** dorsal marker *Dbx1* in NPCs derived from WT and *Tmed2*−/− ESCs overexpressing HA-SMO using the AD protocol. **(E-G)** Immunofluorescence images of **(E)** SMO, and the ER marker ERp72, **(F)** SMO and the Golgi marker RCAS1, and **(G)** SMO and TMED2 in ESCs expressing SMO-HA. Scale = 5μm. The area indicated by the white box is magnified (below). **(H)** TMED2 binds SMO in ESCs. Western analysis of co-immunopurification of endogenous TMED2 and TMED10 with SMO-HA (left) and input (1/25 of IP, right) from extracts of two independent ESCs clones expressing SMO-HA. TMED2 and TMED10 signals in input and IP are from the same exposure. **(I)** Analysis of SMO protein domains for TMED2 binding. Schematic representation of SMO deletions used (CRD, cysteine-rich domain; TM, transmembrane domain; CTD, C-terminal domain). The strength of binding is indicated from pull-down assays in 293T cells shown in Supplementary Fig. 6E. **(J)** SHH treatment disrupts the SMO-TMED2 complex in NPSCs. Western analysis of co-immunopurification of endogenous TMED2 with HA-SMO (top) and input (1/25 of IP, bottom) in NSCs expressing N-term HA tagged SMO. Cells were treated with recombinant SHH for the indicated amount of time. Actin is shown as a loading control. **(K)** Summary of the proposed mechanism for TMED2-regulated SMO secretion from the ER-Golgi compartment.

To assess if SMO-HA at the plasma membrane of *Tmed2* mutant cells was sufficient to activate the HH cascade, we analysed the expression of HH regulated markers in NPCs that were derived from ESCs using the AD protocol. *Olig2*, *Nkx2.2*, *Gli1* and *Ptch1* (Fig. 7B-C, Supplementary Fig. 6B-C) were induced by SHH treatment in control NPCs. In *Tmed2*−/− NPCs, expression of these markers was already increased without SHH treatment. Conversely, expression of the dorsal markers, *Pax7* and *Dbx1*, was strongly decreased (Fig. 7D and Supplementary Fig. 6D). SHH stimulation induced *Olig2*, *Nkx2.2*, *Gli1* and *Ptch1* in *Tmed2*−/− NPCs significantly less efficient than in control cells due to higher basal levels. The effects of the *Tmed2* mutation on elevated HH signalling in the absence of SHH ligand were not observed in NPCs without transgenic SMO-HA (Fig. 5F). Therefore, it is likely that overexpression of SMO-HA contributed to increased basal activity in *Tmed2* mutant NPCs (Fig. 4 and Fig. 5). This observation further suggests that enhanced HH signalling in *Tmed2* mutant embryos also is likely dependent on HH ligand and a *Tmed2* mutation would not lead to ectopic GLI activation.

### *Tmed2* biochemically binds SMO in the ER-Golgi compartment and SHH disrupts the SMO-TMED2 complex

Without HH signal, SMO predominantly localizes in the ER (Incardona et al., 2002). We confirmed ER localization of SMO by immunofluorescence in *Smo−/−* ESCs that carry a *Smo-HA* expression vector (Fig. 7E-F). TMED2 immunofluorescence appeared in an expected Golgi pattern and partially overlapped with SMO-HA in specific and restricted domains (Fig. 7G).

Co-immunoprecipitates (CoIPs) of SMO-HA in ESCs contained TMED2 showing that endogenous TMED2 can bind SMO-HA (Fig. 7H). Notably, TMED10 was not detected in the immunoprecipitate under our conditions suggesting that TMED10 and SMO do not directly interact. To better characterise the interaction between SMO and TMED2, we also tested the binding of SMO variants without the cystein rich domain (CRD) and the C-termininal domain (CTD) in 293T cells (Supplementary Fig. 6E). Pull-down assays indicate that all mutants can bind TMED2. Interestingly, TMED2 binding to the CRD mutant SMO appeared stronger than to wild type SMO (Fig. 7I). The CRD is flexible (Deshpande, Liang et al., 2019, Huang, Zheng et al., 2018) and has been implicated in conformational changes during SMO activation (Rana, Carroll et al., 2013). This observation raised the possibility that TMED2 could contribute to HH signal transduction.

To test if HH signalling regulates the interaction between TMED2 and SMO, we performed HA Co-IPs in NPCs derived from HA-SMO expressing ESCs. TMED2 co-immunoprecipitated with HA-SMO in NPCs (Fig. 7J) consistent with our results in ESCs. Notably, SHH signalling disrupted the interaction between SMO and TMED2 (Fig. 7J). The interaction was lost as early as 1 hour after SHH treatment and remained at low levels in the presence of SHH. The interaction between SMO and TMED2 was also observed with a C-terminally tagged SMO-HA construct (Supplementary Fig. 6F). Also in this case SHH treatment disrupted the SMO-TMED2 complex with similar kinetics. This finding is important as it identifies an early effect downstream of PTCH1 in the activation of the cascade and provides a molecular mechanism for TMED2 function in HH signalling.

## Discussion

Our study identified *Tmed2* and *Tmed10* as members of the p24 family with specific functions in HH signalling. We show that *Tmed2* acts as a novel repressor of HH signals in development by regulating SMO levels at the plasma membrane. Contrary to known negative regulators, such as *Ptch1* and *Sufu*, the mutation of *Tmed2* is not sufficient to induce ectopic GLI activation, but affects the signal strength driven by HH ligands. Importantly we demonstrate that TMED2 specifically interacts with SMO and the SMO-TMED2 complex is disrupted by HH signals. This finding raises the question of how TMED2 could affect SMO abundance at the plasma membrane.

In principle TMED2 could interact with SMO at the cell surface or in endocytic vesicles and be involved in the clearance of the receptor in the absence of HH stimulation. *Atthog* was recently identified as a new regulator of the HH pathway and a mechanism of receptor clearance was proposed (Pusapati et al., 2018). A similar mechanism provides a straight forward explanation for how HH signals could disrupt the SMO-TMED2 complex. HH binding to PTCH1 inhibits cholesterol export from the primary cilium and raising accessibility of cholesterol could disrupt the interaction between SMO and TMED2 by post-translational modifications or conformational changes. However, primary cilia localization of SMO is detected 2 to 4 hours after SHH treatment (Rohatgi et al., 2007) and therefore significantly later than the disruption of the SMO-TMED2 complex, which we observe after 1 hour. In addition, only a minor fraction of TMED2 localizes at the plasma membrane, as detected in our surface biotinylation experiments.

The majority of TMED2 resides in the Golgi, whereas, we detect a tagged SMO construct mainly in the ER compartment consistent with earlier biochemical studies of endogenous SMO (Nachtergaele, Whalen et al., 2013, Rohatgi et al., 2007). Considering SMO and TMED2 localization in ER and Golgi, respectively, an interaction at the interface between these compartments appears likely. Consistent with this idea we detected overlapping staining, which localizes the SMO-TMED2 complex in vesicles that traffic between the ER and Golgi. *Tmed2* mutant cells showed increased levels of SMO at the plasma membrane comparable to control cells after SHH stimulation. In the absence of *Tmed2*, SMO at the plasma membrane had a lower molecular weight suggesting that TMED2 contributes to SMO protein quality control. Retention of immature receptors in the ER-Golgi compartment has previously been implicated as a mechanism of GPCR regulation (Jean-Alphonse & Hanyaloglu, 2011). Our data lead us to propose a model for TMED2 function in the retrograde transport of immature SMO that escaped from the ER compartment (Fig. 7K). In our model, HH stimulation is associated with SMO maturation, and counteracts retention by TMED2. Trafficking of SMO subsequently facilitates GPCR mediated effects at the plasma membrane and regulation of GLI processing at the cilium. Our proposed mechanism for TMED2 function in HH signalling is further supported by a previously proposed role of TMED2 in retention of the PAR-2 receptor (Luo et al., 2007, Luo et al., 2011). However, the finding that SHH disrupts the interaction with SMO raises the question of how SHH signalling could be transduced to the ER-Golgi compartment. Cholesterol abundance at the plasma membrane is sensed in the ER by SCAP/SREBP and regulates cholesterol biosynthesis (Brown, Radhakrishnan et al., 2018). It is tempting to speculate that similar mechanisms could act on SMO to release it form TMED2.

Our screen also identified *Tmed10*. In our hands, TMED10 does not interact directly with SMO. However, TMED10 is required for maintaining TMED2 protein in ESCs, which explains selection of *Tmed10* in our screen. The p24 family plays a crucial role in modulating protein trafficking and signalling strength (Pastor-Cantizano et al., 2016). Specificity of cargo recognition arises from the property of TMED receptors to form multiple homo and hetero dimers. Nonetheless, we expect that in our cellular models multiple factors could be affected by the absence of *Tmed2*. The complexity of defects detected in *Tmed2*−/− embryos supports this view. Differently to *Tmed10*−/− embryos that die before implantation (Denzel, Otto et al., 2000), *Tmed2* mutant embryos develop normally until E7.5 (Jerome-Majewska et al., 2010) and succeed through gastrulation. Therefore, loss of *Tmed2* appears to affect cell-cell interaction mechanisms and signalling mildly in early embryogenesis (Leptin, 2005). *Tmed2* might be specifically important for GPCR regulation. The role of SHH in the modulation of TMED2-SMO interaction provides novel mechanistic insight into the regulatory function of *Tmed2* in HH signalling and development. Further characterization of Golgi-resident proteins identified in our screen will contribute to the understanding of the relevance of the ER and Golgi for HH signalling in the future.

Our study has also defined a, thus far, unrecognized function of SMO in mouse ESCs. Mutation of SMO induces sensitivity to disaggregation. SAG and PMP appear to inactivate the GPCR function of SMO at high concentration and we have used this selection strategy for our screen. Although, the relevance of the role of SMO for preventing cell death in development is unclear at this time, it is consistent with the view that SMO can activate Gi as a GPCR. Notably, the pathway that initiates cell death in the absence of SMO is related to Anoikis in matrix adhesion dependent cells and to dissociation induced cell death of human ESCs (Chen et al., 2010, Ohgushi et al., 2010). In mouse ESCs, the GPCR activity of SMO but not its role in GLI activation is required to confer resistance to dissociation-induced death. SMO antagonizes hyperactivation of ROCK1 and myosin contractility by balancing RAC and ROCK activity in a manner that involves Gi protein activation. HH contributes to embryogenesis through GLI mediated transcription (Briscoe & Therond, 2013). Yet, in mice a *Gli1*/*Gli2*−/− double mutation or primary cilium deficiencies (Murcia, Richards et al., 2000, Park, Bai et al., 2000) show milder phenotypes than a *Shh*/*Ihh* double mutation (Zhang, Ramalho-Santos et al., 2001) and a *Smo* mutation (Zhang et al., 2001). These phenotypic differences suggest additional functions of *Smo.* GLI and cilia independent functions of SMO have been proposed to provide survival signals for neural crest cells (NCCs), where SHH signalling is essential for neural crest migration (Washington Smoak, Byrd et al., 2005). Regulation of SMO at the plasma membrane might be important beyond GLI regulation. Anoikis prevents cells from re-adhering to extracellular matrices and dysplastic growth. Resistance to Anoikis in cancer cells can contribute to the formation of metastasis in distant organs (Paoli, Giannoni et al., 2013). SMO might contribute to tumour dissemination through G-protein mediated mechanisms. Therefore, our observation that classic SMO antagonists are ineffective in blocking SMO induced cell survival in ESCs could be relevant for understanding SMO selective compounds in tumour therapy.

## Acknowledgements

We thank M. de Palma for providing the lentiviral gene trap vector. G. Thinakaran for sharing the TMED10 antibody and H. Roelink for *Smo*−/− ESCs used in preliminary experiments. M. Kruse and M. Torres for helping generating cellular models and reagents. R. Freimann for assistance with cell sorting. M. Ghidinelli, U. Suter, and V. Taylor for help with reagents and discussions. GDM was supported by the ETH Zurich Postdoctoral Fellowship Program as well as the Marie Curie Actions for People COFUND Program. WC and LAJM were supported by a grant from the Natural Sciences and Engineering Research Council of Canada (RGPIN-2015-06699). LJM is a member of the Research Centre of the McGill University Health Centre which is supported in part by FRQS.

This work was supported by grants from the Swiss National Science Foundation (SNF grants 31003A_152814/1 and 31003A_175643/1).

## Author Contributions

GDM and AW designed the study, GDM carried out experiments and performed data analysis. LAJM, WC, and CED prepared mouse embryos. AG performed histological sections and staining. AM contributed to library preparation for NGS sequencing. GDM and AW wrote the manuscript.

## Methods

### Cell culture

ES cells were cultured in chemically defined 2i medium plus LIF as described with minor modifications (Nichols, Jones et al., 2009, Ying et al., 2008). 2i medium was supplemented with non-essential amino acids and 0.35% BSA fraction V. Culture of ES cells on feeders was performed as previously described (Wutz & Jaenisch, 2000). NIH-3T3 and 293T cells were obtained from ATCC and grown in DMEM +10% Fetal bovine serum (FBS) media supplemented with antibiotics. *Ptch1−/−* MEFs were provided by M. P. Scott, Stanford University School of Medicine.

#### Derivation of haploid ESCs

Haploid ESCs from 129S6/SvEvTac Oocytes (ha129DM1) were derived as previously described (Leeb & Wutz, 2011). Briefly, oocytes were isolated from superovulated female mice and activated in KSOM medium using 5 mM strontium chloride and 2 mM EGTA. Embryos were subsequently cultured in Cleavage (Cook Medical, G20720) medium microdrops covered by mineral oil. After 4-5 days embryos at the morula or blastocyst stage were treated with acidic Tyrode’s solution to remove the zona pellucida and ESCs derivation was performed as described previously.

### Mice and embryo section

All animal procedures were approved by the veterinary office of the Canton of Zurich and the Canadian Council on Animal Research. The *Tmed2^99/99J^* (−/−) mouse line was described previously (Jerome-Majewska et al., 2010) and genotyped by PCR using the following primers: *Tmed2*In4F (AAGTGCACAGCTGAGTGGT) and *Tmed2In4R* (CACAGTGTCTGACCCCCTTT). Embryos of E10.5 *Tmed2*−/− mice were compared to E9.0 WT embryos (CD1 strain). For sections, embryos were fixed with 4% paraformaldehyde (wt/vol), cryoprotected with 30% sucrose (wt/vol) overnight, and embedded in OCT (Leica, Germany). 10μm horizontal cryotome sections (E9.0, E9.5 and E10.5) were processed for immunofluorescence staining.

### Reagents

#### Growth factors and chemicals

The following growth factors and chemicals were used: Leukemia inhibitory factor (LIF, recombinant purified as GST-fusion protein, homemade), mouse sonic hedgehog C24II (SHH, recombinant purified as GST-fusion protein, homemade), mouse Epidermal growth factor (EGF, PeproTech), human basic Fibroblast growth factor (bFGF, PeproTech), CHIR99021 (Axon Medchem), PD0325901 (Axon Medchem), Smoothened agonist (SAG,Calbiochem), Purmorphamine (PMP, Calbiochem), Cyclopamine-KAAD (Calbiochem), SANT-1 (Sigma), Nocodazole (Sigma).

#### SHH production

The pcDNA3-Shh (N) (Addgene, #37680) was used as template to amplify the cDNA coding residues 24-197 of mouse *Shh*. By PCR a Cys24-Ile-Ile substitution and a Factor Xa cleavage site were introduced. The construct was cloned in the pGEX vector and transformed in BL21 (DE3) pLysS *E.coli* cells. The transformed cultures were grown in a shaking incubator at 37°C until OD600 of 0.8 was reached, at which point the temperature was switched to 30°C and the expression was induced with 0.5 mM IPTG. After 4 hrs cells were collected by centrifugation, frozen and stored at −80°C until the day of experiment. ShhNC24II-expressing *E.coli* pellets (0.1 L culture) were thawed, resuspended in 5 ml of lysis buffer (25 mM sodium phosphate (pH 8.0), 150 mM NaCl, 1 mM EDTA, 0.5 mM DTT and 1 mM PMSF), disrupted by sonication, and cleared by centrifugation for 30 min at 25000 x g. The supernatant was added to Glutathione Sepharose 4B Resin (Sigma, GE17-0756-01) and incubated O/N at 4°C. The resin was collected and washed 5 times in Lysis Buffer. The protein was eluted by digestion with Factor Xa protease (Sigma). Factor Xa protease was removed using p-Aminobenzamidine-agarose (Sigma) and recombinant SHH was purified with Detoxi-Gel™ Endotoxin Removing Gel (ThermoFisher) and dialyzed. SHH was supplemented with 10% glycerol, aliquoted into 100 μl batches, flash-frozen in liquid nitrogen and stored at −80°C.

#### Antibodies

The following antibodies were used: αTMED2 (Santa Cruz Biotechnology, sc-376459), αTMED10 (sc-137003), αSMO (sc-166685), αActin (Sigma A5316), αHA (Roche 12013819001), αARL13B (Proteintech 17711-1-AP), αRCAS1 (Cell Signaling #12290), αERp72 (Cell Signaling #5033), αOLIG2 (Merck-Millipore AB9610), αNKX2.2 (DSHB), αNKX6.1 (DSHB), αPAX7 (DSHB), αPAX6 (Novus NBP195459).

### Cell treatment

#### Flow cytometry and activated cell sorting

For derivation and maintenance of haploid ESCs, cell sorting for DNA content was performed after staining with 15 mg/ml Hoechst 33342 (Invitrogen) on a MoFlo flow sorter (Beckman Coulter) selecting for the haploid 1n peak.

#### Transfection and stable cell line generation

For generation of *Tmed2*−/−, *Tmed10*−/−, and *Smo*−/− ESCs the following gRNAs sequences were cloned in the pX458_pSpCas9(BB)-2A-GFP vector (Addgene, #48138):

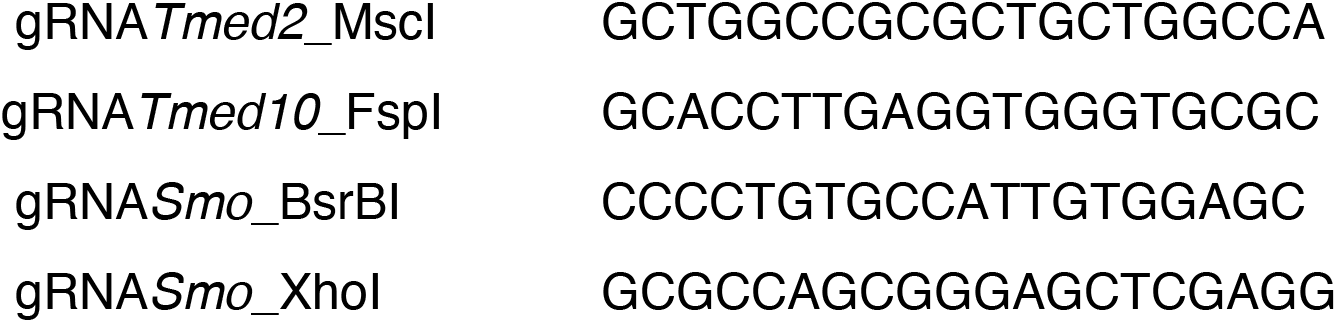

Briefly, plasmid vectors were used for transfection (Lipofecatmine2000) of haploid 129DM1, 48 hr later, cells were sorted for green fluorescence and plated at low density for isolating individual clones. Mutations were identified by PCR on genomic DNA and sequencing using the following primers:

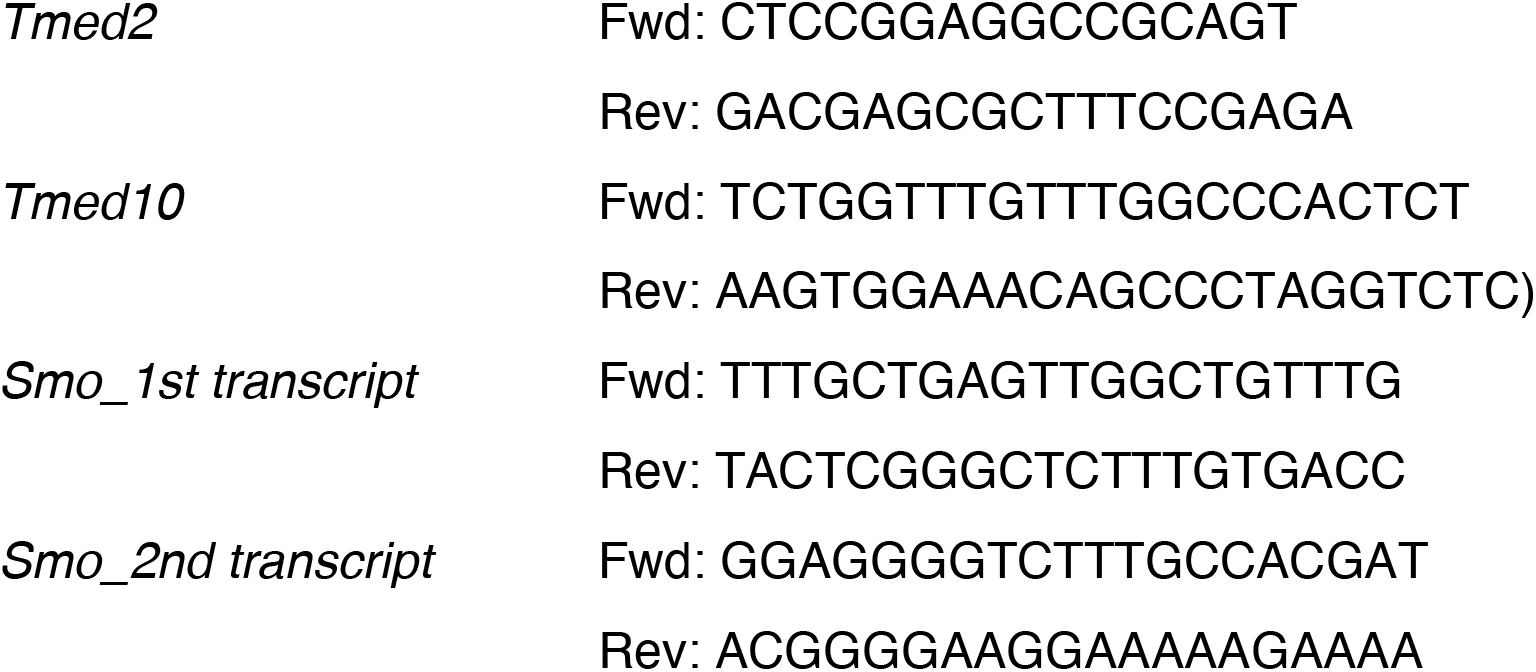

For derivation of HA-*Smo* and *Smo*-HA ESCs mouse *Smo* was amplified by PCR from the pGEN-*mSmo* (Addgene, #37673) and introduced in the PB-HA-IRES-Neo vector. Stable integration of the construct was obtained in *Smo*−/− ESCs by co-transfection of the Piggybac and the PBase plasmids. Cells were plated by limiting dilution and integration events selected with G418. Two clones characterized by high and low HA-SMO or SMO-HA expression were expanded. *Smo*-HA *Tmed2*−/− ESCs were derived from *Smo*-HA ESCs with low SMO-HA expression as previously described.

Transfections of NIH-3T3 cells with siRNA targeting mouse *Tmed2* (Fwd: CACCUCUAAUUGAAUUGAACAAGCA, Rev: UGCUUGUUCAAUUCAAUUAGAGGUGAU) were performed with RNAiMax (Invitrogen). The Negative Control DsiRNA (IDT, 73481795) was used as transfection control.

#### Drug treatment and survival assay

ESCs and NIH-3T3 were treated with SHH (50-500 ng/ml), SAG (0.5-5 μM), PMP (1-10 μM), Cyclopamine-KAAD (1-5 μM) or SANT-1 (10-50 μM). After 48 hours cells were trypsinized and counted. Survival is expressed normalizing cell counts to DMSO treated sample. To evaluate effects of compounds on GLI transcriptional activity cells were treated with SHH (500 ng/ml), SAG (5 μM), PMP (10 μM) or Cyclopamine-KAAD (5 μM) or SANT-1 (50 μM).

#### Hedgehog signaling assays

NIH-3T3 cells were grown to confluency in DMEM containing 10% FBS. Confluent cells were cultured in 0.5% FBS DMEM for 24 hr to allow ciliogenesis prior to treatment with drugs and/or ligands in DMEM containing 0.5% FBS for various times, as indicated in the figures.

#### NPCs derivation from ESCs

Differentiation of ESCs to NPCs in 2D conditions was previously described (Kutejova, Sasai et al., 2016). Briefly, the cells were plated on Matrigel (Matrigel® hESC-Qualified Matrix, LDEV-free (Corning)) in N2B27 media (DMEM-F12 (Gibco) and Neurobasal Medium (Gibco) mixed in 1:1 ratio and supplemented with N-2 supplement, B-27 supplement, 1% penicillin/streptomycin), 2 mM L-glutamine, 40 mg/ml Bovine Serum Albumin, and 55 mM 2-mercaptoethanol). On day 0 and day 1, cells were cultured in N2B27 with 10 ng/ml bFGF. On day 2, the media was changed and the cells were cultured in N2B27 with 10 ng/ml bFGF and 5 mM CHIR9902. On day 3, the media was changed and the cells were cultured in N2B27 supplemented with retinoic acid (RA, 100nM), and SHH (1-4μg/ml). On day 4, an equal volume of N2B27 with 100 nM RA was added to each well diluting each treatment condition by half. On day 5 the cells were processed for further analysis. For derivation of neuralized EBs, ESCs were plated in EBs media (DMEM-F12 (Gibco) and Neurobasal Medium (Gibco) (1:1 ratio) supplemented with 200mM L-glutamine and 10% of Knockout serum replacement) on Sphericalplate 5D dishes (Kugelmeiers AG). On day 2, EBs were transferred in 10cm tissue culture dishes and treated with RA (10nM). EBs were treated with SHH (4μg/ml) the day after and neuralized EBs were collected and processed for analysis at day 5.

### Genome-wide screening

Haploid ESCs were mutagenized using a lentivirus system with a gene-trap cassette (De Palma, Montini et al., 2005). The pRRLsin-PPT-SA-PuroGFP-Wpre-pA plasmid was transfected with lentiviral packaging plasmids in 293T. Virus was collected and concentrated by ultracentrifugation to obtain a high viral titer. 6×10^7^ sorted haploid ESCs were infected with virus and plated on 145 cm^2^ dishes pre-coated with MEFs. After 2 days, cells were collected and split into control and selected sample. Control cells were directly lysed and DNA was extracted. For selection, PMP was added to the media at a concentration of 10 μM. Every 2 days, cells were passaged until day 14. Cells not infected were grown in parallel and treated like infected cells. For identification of genomic insertion sites of genetrap viruses next generation sequencing libraries were prepared and sequenced on an Illumina MiSeq device as described (Monfort, Di Minin et al., 2018). In brief, genomic sequences adjacent to the end of the viral LTR were enriched by linear amplification PCR (LAMPCR) using a high-fidelity DNA polymerase (Invitrogen, #12346094) with a biotinylated primer targeting the viral genome. Single stranded LAM PCR products were captured on Streptavidin coated magnetic beads (Dynabeads M270, Invitrogen #65305). Subsequently, an oligonucleotide containing the Illumina P7 adaptor sequence was ligated to the 3’ end of the single stranded LAM PCR fragments using Circligase (Epicentre/Illumina, #CL9025K). A P5 adaptor was added by PCR. The resulting DNA was purified and concentrated using the MinElute PCR Purification Kit (Qiagen, #28004) before loading it on the Illumina MiSeq flow cell. Sequencing was performed using 75 cycles, pair-end runs on an Illumina MiSeq sequencer using MiSeq v3 kits (Illumina, #MS1023001). Computational analysis of NGS datasets was performed using the HaSAPPy package (Di Minin et al., 2018).

### RNA isolation and qPCR

Total RNA was isolated from ESCs using the QiAshredder (Qiagen, #79656) and purified using RNeasy Mini Kit (Qiagen, #74104) and on-column DNase I digestion (Qiagen, #79254). cDNA for real time PCR was synthesized using the QuantiTect Reverse Transcription Kit (Quiagen, #205313). Realtime quantitative PCR reactions were performed using SYBR Green and a LightCycler 480 System (Roche). Relative expression of the target gene was normalized to *Eif4a2* and *Sdha* expression levels. Primer sequences:

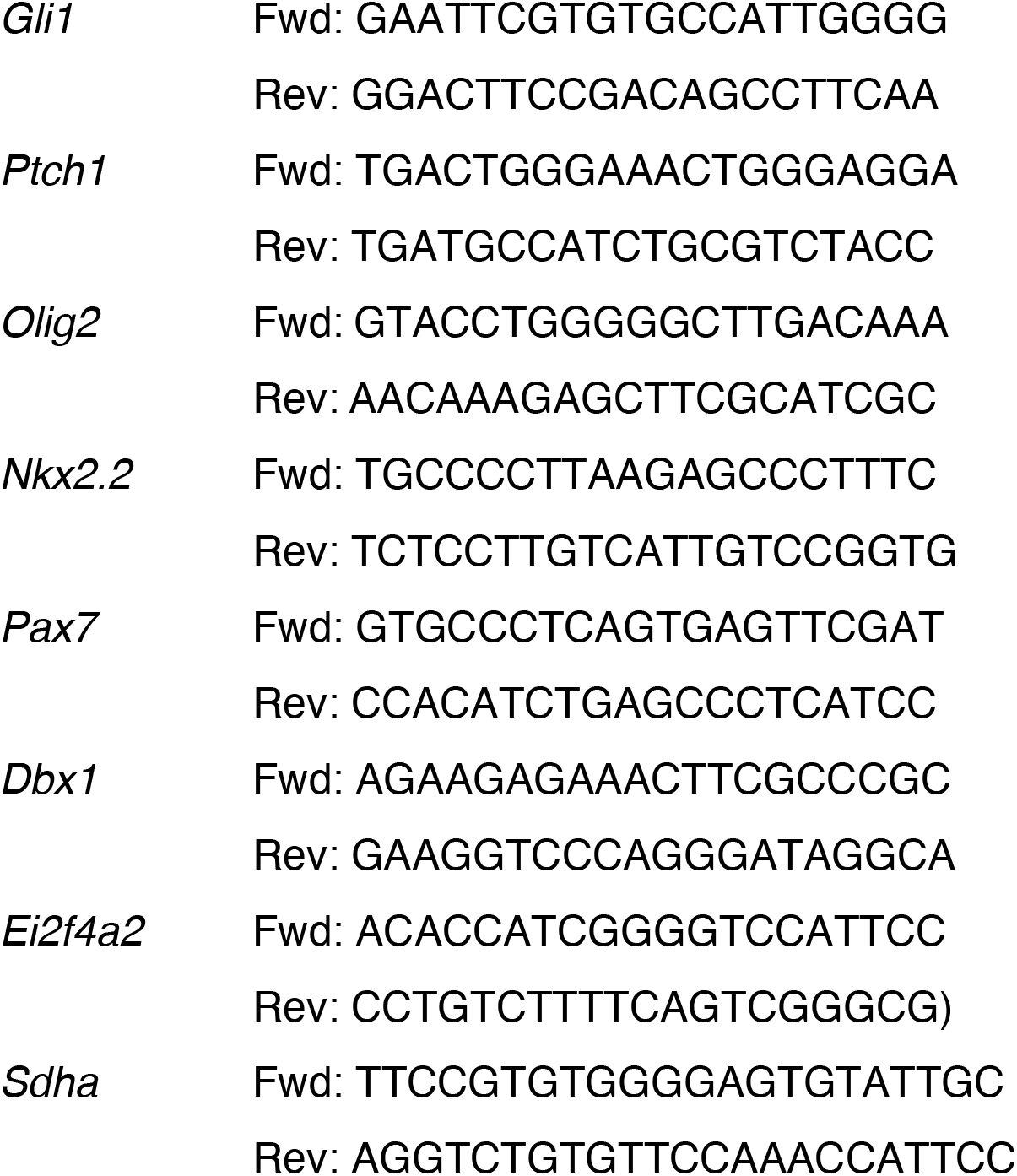

### Protein extraction and Immunoblotting

Whole cell extracts from NIH-3T3, ES cells, or NPCs were prepared in RIPA lysis buffer (50 mM Tris-HCl pH-7.4, 150 mM NaCl, 2% NP-40, 0.25% Deoxycholate, 0.1% SDS, 1mM DTT, 10% glycerol, protease inhibitors). Samples were resuspended in Leammli sample buffer, denatured, and subjected to SDS-PAGE. The resolved proteins were transferred onto a nitrocellulose membrane (Bio-Rad) using a wet electroblotting system (Bio-Rad) followed by immunoblotting.

#### Co-immunoprecipitation experiments

For Co-IP experiments cells were lysed in the Membrane lysis buffer (50mM Tris-HCl pH 7.5, 1mM β-mercaptoethanol, 150mM NaCl, 1% ChAPS and proteases inhibitors) and quantified. An equal amount of proteins was incubated with anti-HA affinity matrix (Sigma, #11815016001) to pull-down SMO-HA. After 1h samples were washed, denatured in Laemmli sample buffer and urea, and subjected to SDS-PAGE.

#### Plasma membrane protein purification

Biotinylation of cell surface proteins was performed as described previously (Karhemo, Ravela et al., 2012). Briefly, ESCs were differentiated to NPCs as previously described in 10cm dishes. Cells were incubated for 30 min with Sulfo-NHS-SS-Biotin (Covachem, #14207) on ice and lysed in Membrane lysis buffer. Lysates were quantified and an equal amount of protein lysate was incubated on Pierce™ High Capacity Streptavidin Agarose (Pierce, # 20357) for 1h. Samples were washed, denatured in Laemmli sample buffer and urea and subjected to SDS-PAGE.

### Immunofluorescence and localization studies

Cells were fixed in 4% PFA for 15 min at 37°C and washed in PBS. Cells were incubated in blocking buffer (10% donkey serum, 0.3% triton, 1x PBS) for 30 min. Primary antibodies diluted in antibody buffer (1% BSA, 0.3% triton, 1x PBS) were added for 1h or O/N. Secondary antibodies and DAPI were added for 45 min in antibody buffer. To improve detection of TMED2 protein, cells were incubated in Golgi buffer (0.5% SDS, 1% 2-Mercaptoethanol, 10% donkey serum in PBS) for 30 min at 60°C before incubation with the primary antibody. Samples were viewed on a Zeiss Image Z1 microscope equipped with an X-Cite 120 illuminator (EXFO) and a Leica SP8 confocal microscope.

For immunofluorescence staining of embryo sections, the sections were subjected to the following antigen retrieval procedure to increase signaling. Samples were incubated in H_2_O_2_ 0.3% diluted in phosphate-buffered saline (PBS) for 30’. Antigens were recovered at 105 °C with pressure cooker device for 15 minutes in pre-warmed sodium citrate solution (1.8 mM citric acid monohydrate, 8.2 mM sodium citrate tribasic dihydrate). Sections were cooled down to room temperature for 2 hours and then incubated in HCl 2N for 3’ at 37°C. Sections were incubated in TNB blocking solution (0.1 M Tris-HCl pH 7.5, 0.15 M NaCl, 0.5% Blocking reagent PerkinElmer_FP1012) at room temperature for 30’. Primary and secondary staining were performed as previously described.

### Quantification and statistical analysis

Statistical analysis was performed using Python and GraphPad Prism. Data are presented as mean-centered and the standard deviation. All experiments were repeated with at least three independent biological replicates, unless otherwise stated.

### Data availability

The sequencing datasets are deposited in the NCBI Short-Read Archive (http://www.ncbi.nlm.nih.gov/sra) and can be accessed using accession numbers SRR8449980 and SRR8449981.

### Code availability

All custom scripts used in this study are available from the corresponding author on reasonable request.

## References

Ast T, Cohen G, Schuldiner M (2013) A network of cytosolic factors targets SRP-independent proteins to the endoplasmic reticulum. Cell 152: 1134–45

Bijlsma MF, Damhofer H, Roelink H (2012) Hedgehog-stimulated chemotaxis is mediated by smoothened located outside the primary cilium. Sci Signal 5: ra60

Brandizzi F, Barlowe C (2013) Organization of the ER-Golgi interface for membrane traffic control. Nat Rev Mol Cell Biol 14: 382–92

Brennan D, Chen X, Cheng L, Mahoney M, Riobo NA (2012) Noncanonical Hedgehog signaling. Vitam Horm 88: 55–72

Breslow DK, Hoogendoorn S, Kopp AR, Morgens DW, Vu BK, Kennedy MC, Han K, Li A, Hess GT, Bassik MC, Chen JK, Nachury MV (2018) A CRISPR-based screen for Hedgehog signaling provides insights into ciliary function and ciliopathies. Nat Genet 50: 460–471

Briscoe J, Therond PP (2013) The mechanisms of Hedgehog signalling and its roles in development and disease. Nat Rev Mol Cell Biol 14: 416–29

Brown MS, Radhakrishnan A, Goldstein JL (2018) Retrospective on Cholesterol Homeostasis: The Central Role of Scap. Annu Rev Biochem 87: 783–807

Bryce NS, Schevzov G, Ferguson V, Percival JM, Lin JJ, Matsumura F, Bamburg JR, Jeffrey PL, Hardeman EC, Gunning P, Weinberger RP (2003) Specification of actin filament function and molecular composition by tropomyosin isoforms. Mol Biol Cell 14: 1002–16

Byrne EFX, Sircar R, Miller PS, Hedger G, Luchetti G, Nachtergaele S, Tully MD, Mydock-McGrane L, Covey DF, Rambo RP, Sansom MSP, Newstead S, Rohatgi R, Siebold C (2016) Structural basis of Smoothened regulation by its extracellular domains. Nature 535: 517–522

Chang HM, Martinez NJ, Thornton JE, Hagan JP, Nguyen KD, Gregory RI (2012) Trim71 cooperates with microRNAs to repress Cdkn1a expression and promote embryonic stem cell proliferation. Nat Commun 3: 923

Chen G, Hou Z, Gulbranson DR, Thomson JA (2010) Actin-myosin contractility is responsible for the reduced viability of dissociated human embryonic stem cells. Cell Stem Cell 7: 240–8

Chen JK, Taipale J, Young KE, Maiti T, Beachy PA (2002) Small molecule modulation of Smoothened activity. Proc Natl Acad Sci U S A 99: 14071–6

Cooper MK, Wassif CA, Krakowiak PA, Taipale J, Gong R, Kelley RI, Porter FD, Beachy PA (2003) A defective response to Hedgehog signaling in disorders of cholesterol biosynthesis. Nat Genet 33: 508–13

Corbit KC, Aanstad P, Singla V, Norman AR, Stainier DY, Reiter JF (2005) Vertebrate Smoothened functions at the primary cilium. Nature 437: 1018–21

Dai J, Ma J, Yu B, Zhu Z, Hu Y (2018) Long Noncoding RNA TUNAR Represses Growth, Migration, and Invasion of Human Glioma Cells Through Regulating miR-200a and Rac1. Oncol Res 27: 107–115

De Palma M, Montini E, Santoni de Sio FR, Benedicenti F, Gentile A, Medico E, Naldini L (2005) Promoter trapping reveals significant differences in integration site selection between MLV and HIV vectors in primary hematopoietic cells. Blood 105: 2307–15

Denef N, Neubuser D, Perez L, Cohen SM (2000) Hedgehog induces opposite changes in turnover and subcellular localization of patched and smoothened. Cell 102: 521–31

Denzel A, Otto F, Girod A, Pepperkok R, Watson R, Rosewell I, Bergeron JJ, Solari RC, Owen MJ (2000) The p24 family member p23 is required for early embryonic development. Curr Biol 10: 55–8

Deshpande I, Liang J, Hedeen D, Roberts KJ, Zhang Y, Ha B, Latorraca NR, Faust B, Dror RO, Beachy PA, Myers BR, Manglik A (2019) Smoothened stimulation by membrane sterols drives Hedgehog pathway activity. Nature 571: 284–288

Desouza-Armstrong M, Gunning PW, Stehn JR (2017) Tumor suppressor tropomyosin Tpm2.1 regulates sensitivity to apoptosis beyond anoikis characterized by changes in the levels of intrinsic apoptosis proteins. Cytoskeleton (Hoboken) 74: 233–248

Dessaud E, McMahon AP, Briscoe J (2008) Pattern formation in the vertebrate neural tube: a sonic hedgehog morphogen-regulated transcriptional network. Development 135: 2489–503

Di Minin G, Postlmayr A, Wutz A (2018) HaSAPPy: A tool for candidate identification in pooled forward genetic screens of haploid mammalian cells. PLoS Comput Biol 14: e1005950

Dong C, Filipeanu CM, Duvernay MT, Wu G (2007) Regulation of G protein-coupled receptor export trafficking. Biochim Biophys Acta 1768: 853–70

Fan CW, Chen B, Franco I, Lu J, Shi H, Wei S, Wang C, Wu X, Tang W, Roth MG, Williams NS, Hirsch E, Chen C, Lum L (2014) The Hedgehog pathway effector smoothened exhibits signaling competency in the absence of ciliary accumulation. Chem Biol 21: 1680–9

Frank-Kamenetsky M, Zhang XM, Bottega S, Guicherit O, Wichterle H, Dudek H, Bumcrot D, Wang FY, Jones S, Shulok J, Rubin LL, Porter JA (2002) Small-molecule modulators of Hedgehog signaling: identification and characterization of Smoothened agonists and antagonists. J Biol 1: 10

Girardi D, Barrichello A, Fernandes G, Pereira A (2019) Targeting the Hedgehog Pathway in Cancer: Current Evidence and Future Perspectives. Cells 8

Gong X, Qian H, Cao P, Zhao X, Zhou Q, Lei J, Yan N (2018) Structural basis for the recognition of Sonic Hedgehog by human Patched1. Science 361

Haystead TA (2005) ZIP kinase, a key regulator of myosin protein phosphatase 1. Cell Signal 17: 1313–22

Hou W, Gupta S, Beauchamp MC, Yuan L, Jerome-Majewska LA (2017) Non-alcoholic fatty liver disease in mice with heterozygous mutation in TMED2. PLoS One 12: e0182995

Hou W, Jerome-Majewska LA (2018) TMED2/emp24 is required in both the chorion and the allantois for placental labyrinth layer development. Dev Biol 444: 20–32

Huang P, Nedelcu D, Watanabe M, Jao C, Kim Y, Liu J, Salic A (2016) Cellular Cholesterol Directly Activates Smoothened in Hedgehog Signaling. Cell 166: 1176–1187 e14

Huang P, Zheng S, Wierbowski BM, Kim Y, Nedelcu D, Aravena L, Liu J, Kruse AC, Salic A (2018) Structural Basis of Smoothened Activation in Hedgehog Signaling. Cell 175: 295–297

Incardona JP, Gruenberg J, Roelink H (2002) Sonic hedgehog induces the segregation of patched and smoothened in endosomes. Curr Biol 12: 983–95

Jacob LS, Wu X, Dodge ME, Fan CW, Kulak O, Chen B, Tang W, Wang B, Amatruda JF, Lum L (2011) Genome-wide RNAi screen reveals disease-associated genes that are common to Hedgehog and Wnt signaling. Sci Signal 4: ra4

Jean-Alphonse F, Hanyaloglu AC (2011) Regulation of GPCR signal networks via membrane trafficking. Mol Cell Endocrinol 331: 205–14

Jerome-Majewska LA, Achkar T, Luo L, Lupu F, Lacy E (2010) The trafficking protein Tmed2/p24beta(1) is required for morphogenesis of the mouse embryo and placenta. Dev Biol 341: 154–66

Joo EE, Yamada KM (2014) MYPT1 regulates contractility and microtubule acetylation to modulate integrin adhesions and matrix assembly. Nat Commun 5: 3510

Karhemo PR, Ravela S, Laakso M, Ritamo I, Tatti O, Makinen S, Goodison S, Stenman UH, Holtta E, Hautaniemi S, Valmu L, Lehti K, Laakkonen P (2012) An optimized isolation of biotinylated cell surface proteins reveals novel players in cancer metastasis. J Proteomics 77: 87–100

Kinnebrew M, Iverson EJ, Patel BB, Pusapati GV, Kong JH, Johnson KA, Luchetti G, Eckert KM, McDonald JG, Covey DF, Siebold C, Radhakrishnan A, Rohatgi R (2019) Cholesterol accessibility at the ciliary membrane controls hedgehog signaling. Elife 8

Kong JH, Siebold C, Rohatgi R (2019) Biochemical mechanisms of vertebrate hedgehog signaling. Development 146

Kowatsch C, Woolley RE, Kinnebrew M, Rohatgi R, Siebold C (2019) Structures of vertebrate Patched and Smoothened reveal intimate links between cholesterol and Hedgehog signalling. Curr Opin Struct Biol 57: 204–214

Kutejova E, Sasai N, Shah A, Gouti M, Briscoe J (2016) Neural Progenitors Adopt Specific Identities by Directly Repressing All Alternative Progenitor Transcriptional Programs. Dev Cell 36: 639–53

Leeb M, Dietmann S, Paramor M, Niwa H, Smith A (2014) Genetic exploration of the exit from self-renewal using haploid embryonic stem cells. Cell Stem Cell 14: 385–93

Leeb M, Wutz A (2011) Derivation of haploid embryonic stem cells from mouse embryos. Nature 479: 131–4

Leptin M (2005) Gastrulation movements: the logic and the nuts and bolts. Dev Cell 8: 305–20

Lindsay AJ, McCaffrey MW (2015) Rab antibody characterization: comparison of Rab14 antibodies. Methods Mol Biol 1298: 161–71

Luchetti G, Sircar R, Kong JH, Nachtergaele S, Sagner A, Byrne EF, Covey DF, Siebold C, Rohatgi R (2016) Cholesterol activates the G-protein coupled receptor Smoothened to promote Hedgehog signaling. Elife 5

Luo W, Wang Y, Reiser G (2007) p24A, a type I transmembrane protein, controls ARF1-dependent resensitization of protease-activated receptor-2 by influence on receptor trafficking. J Biol Chem 282: 30246–55

Luo W, Wang Y, Reiser G (2011) Proteinase-activated receptors, nucleotide P2Y receptors, and mu-opioid receptor-1B are under the control of the type I transmembrane proteins p23 and p24A in post-Golgi trafficking. J Neurochem 117: 71–81

Milenkovic L, Scott MP, Rohatgi R (2009) Lateral transport of Smoothened from the plasma membrane to the membrane of the cilium. J Cell Biol 187: 365–74

Monfort A, Di Minin G, Postlmayr A, Freimann R, Arieti F, Thore S, Wutz A (2015) Identification of Spen as a Crucial Factor for Xist Function through Forward Genetic Screening in Haploid Embryonic Stem Cells. Cell Rep 12: 554–61

Monfort A, Di Minin G, Wutz A (2018) Screening for Factors Involved in X Chromosome Inactivation Using Haploid ESCs. Methods Mol Biol 1861: 1–18

Moretti F, Bergman P, Dodgson S, Marcellin D, Claerr I, Goodwin JM, DeJesus R, Kang Z, Antczak C, Begue D, Bonenfant D, Graff A, Genoud C, Reece-Hoyes JS, Russ C, Yang Z, Hoffman GR, Mueller M, Murphy LO, Xavier RJ et al. (2018) TMEM41B is a novel regulator of autophagy and lipid mobilization. EMBO Rep 19

Murcia NS, Richards WG, Yoder BK, Mucenski ML, Dunlap JR, Woychik RP (2000) The Oak Ridge Polycystic Kidney (orpk) disease gene is required for left-right axis determination. Development 127: 2347–55

Myers BR, Neahring L, Zhang Y, Roberts KJ, Beachy PA (2017) Rapid, direct activity assays for Smoothened reveal Hedgehog pathway regulation by membrane cholesterol and extracellular sodium. Proc Natl Acad Sci U S A 114: E11141–E11150

Nachtergaele S, Whalen DM, Mydock LK, Zhao Z, Malinauskas T, Krishnan K, Ingham PW, Covey DF, Siebold C, Rohatgi R (2013) Structure and function of the Smoothened extracellular domain in vertebrate Hedgehog signaling. Elife 2: e01340

Nichols J, Jones K, Phillips JM, Newland SA, Roode M, Mansfield W, Smith A, Cooke A (2009) Validated germline-competent embryonic stem cell lines from nonobese diabetic mice. Nat Med 15: 814–8

Niewiadomski P, Niedziolka SM, Markiewicz L, Uspienski T, Baran B, Chojnowska K (2019) Gli Proteins: Regulation in Development and Cancer. Cells 8

Ogden SK, Fei DL, Schilling NS, Ahmed YF, Hwa J, Robbins DJ (2008) G protein Galphai functions immediately downstream of Smoothened in Hedgehog signalling. Nature 456: 967–70

Ohgushi M, Matsumura M, Eiraku M, Murakami K, Aramaki T, Nishiyama A, Muguruma K, Nakano T, Suga H, Ueno M, Ishizaki T, Suemori H, Narumiya S, Niwa H, Sasai Y (2010) Molecular pathway and cell state responsible for dissociation-induced apoptosis in human pluripotent stem cells. Cell Stem Cell 7: 225–39

Pak E, Segal RA (2016) Hedgehog Signal Transduction: Key Players, Oncogenic Drivers, and Cancer Therapy. Dev Cell 38: 333–44

Pandit T, Ogden SK (2017) Contributions of Noncanonical Smoothened Signaling During Embryonic Development. J Dev Biol 5

Paoli P, Giannoni E, Chiarugi P (2013) Anoikis molecular pathways and its role in cancer progression. Biochim Biophys Acta 1833: 3481–3498

Park HL, Bai C, Platt KA, Matise MP, Beeghly A, Hui CC, Nakashima M, Joyner AL (2000) Mouse Gli1 mutants are viable but have defects in SHH signaling in combination with a Gli2 mutation. Development 127: 1593–605

Pastor-Cantizano N, Montesinos JC, Bernat-Silvestre C, Marcote MJ, Aniento F (2016) p24 family proteins: key players in the regulation of trafficking along the secretory pathway. Protoplasma 253: 967–85

Peng T, Frank DB, Kadzik RS, Morley MP, Rathi KS, Wang T, Zhou S, Cheng L, Lu MM, Morrisey EE (2015) Hedgehog actively maintains adult lung quiescence and regulates repair and regeneration. Nature 526: 578–82

Petrov K, Wierbowski BM, Salic A (2017) Sending and Receiving Hedgehog Signals. Annu Rev Cell Dev Biol 33: 145–168

Petrova R, Joyner AL (2014) Roles for Hedgehog signaling in adult organ homeostasis and repair. Development 141: 3445–57

Phillips SE, Ile KE, Boukhelifa M, Huijbregts RP, Bankaitis VA (2006) Specific and nonspecific membrane-binding determinants cooperate in targeting phosphatidylinositol transfer protein beta-isoform to the mammalian trans-Golgi network. Mol Biol Cell 17: 2498–512

Polizio AH, Chinchilla P, Chen X, Kim S, Manning DR, Riobo NA (2011a) Heterotrimeric Gi proteins link Hedgehog signaling to activation of Rho small GTPases to promote fibroblast migration. J Biol Chem 286: 19589–96

Polizio AH, Chinchilla P, Chen X, Manning DR, Riobo NA (2011b) Sonic Hedgehog activates the GTPases Rac1 and RhoA in a Gli-independent manner through coupling of smoothened to Gi proteins. Sci Signal 4: pt7

Pusapati GV, Kong JH, Patel BB, Krishnan A, Sagner A, Kinnebrew M, Briscoe J, Aravind L, Rohatgi R (2018) CRISPR Screens Uncover Genes that Regulate Target Cell Sensitivity to the Morphogen Sonic Hedgehog. Dev Cell 44: 113–129 e8

Qi C, Di Minin G, Vercellino I, Wutz A, Korkhov VM (2019) Structural basis of sterol recognition by human hedgehog receptor PTCH1. Sci Adv 5: eaaw6490

Qi X, Schmiege P, Coutavas E, Li X (2018a) Two Patched molecules engage distinct sites on Hedgehog yielding a signaling-competent complex. Science 362

Qi X, Schmiege P, Coutavas E, Wang J, Li X (2018b) Structures of human Patched and its complex with native palmitoylated sonic hedgehog. Nature 560: 128–132

Qian H, Cao P, Hu M, Gao S, Yan N, Gong X (2019) Inhibition of tetrameric Patched1 by Sonic Hedgehog through an asymmetric paradigm. Nat Commun 10: 2320

Rana R, Carroll CE, Lee HJ, Bao J, Marada S, Grace CR, Guibao CD, Ogden SK, Zheng JJ (2013) Structural insights into the role of the Smoothened cysteine-rich domain in Hedgehog signalling. Nat Commun 4: 2965

Riobo NA, Saucy B, Dilizio C, Manning DR (2006) Activation of heterotrimeric G proteins by Smoothened. Proc Natl Acad Sci U S A 103: 12607–12

Rohatgi R, Milenkovic L, Scott MP (2007) Patched1 regulates hedgehog signaling at the primary cilium. Science 317: 372–6

Shen F, Cheng L, Douglas AE, Riobo NA, Manning DR (2013) Smoothened is a fully competent activator of the heterotrimeric G protein G(i). Mol Pharmacol 83: 691–7

Shi J, Wu X, Surma M, Vemula S, Zhang L, Yang Y, Kapur R, Wei L (2013) Distinct roles for ROCK1 and ROCK2 in the regulation of cell detachment. Cell Death Dis 4: e483

Solimini NL, Liang AC, Xu C, Pavlova NN, Xu Q, Davoli T, Li MZ, Wong KK, Elledge SJ (2013) STOP gene Phactr4 is a tumor suppressor. Proc Natl Acad Sci U S A 110: E407–14

Stepanchick A, Breitwieser GE (2010) The cargo receptor p24A facilitates calcium sensing receptor maturation and stabilization in the early secretory pathway. Biochem Biophys Res Commun 395: 136–40

Strating JR, Martens GJ (2009) The p24 family and selective transport processes at the ER-Golgi interface. Biol Cell 101: 495–509

Sun MS, Zhang J, Jiang LQ, Pan YX, Tan JY, Yu F, Guo L, Yin L, Shen C, Shu HB, Liu Y (2018) TMED2 Potentiates Cellular IFN Responses to DNA Viruses by Reinforcing MITA Dimerization and Facilitating Its Trafficking. Cell Rep 25: 3086–3098 e3

Tabara LC, Vicente JJ, Biazik J, Eskelinen EL, Vincent O, Escalante R (2018) Vacuole membrane protein 1 marks endoplasmic reticulum subdomains enriched in phospholipid synthesizing enzymes and is required for phosphoinositide distribution. Traffic 19: 624–638

Takahashi S, Lee J, Kohda T, Matsuzawa A, Kawasumi M, Kanai-Azuma M, Kaneko-Ishino T, Ishino F (2014) Induction of the G2/M transition stabilizes haploid embryonic stem cells. Development 141: 3842–7

Takubo T, Wakui S, Daigo K, Kurokata K, Ohashi T, Katayama K, Hino M (2003) Expression of non-muscle type myosin heavy polypeptide 9 (MYH9) in mammalian cells. Eur J Histochem 47: 345–52

Teperino R, Amann S, Bayer M, McGee SL, Loipetzberger A, Connor T, Jaeger C, Kammerer B, Winter L, Wiche G, Dalgaard K, Selvaraj M, Gaster M, Lee-Young RS, Febbraio MA, Knauf C, Cani PD, Aberger F, Penninger JM, Pospisilik JA et al. (2012) Hedgehog partial agonism drives Warburg-like metabolism in muscle and brown fat. Cell 151: 414–26

Vetrivel KS, Gong P, Bowen JW, Cheng H, Chen Y, Carter M, Nguyen PD, Placanica L, Wieland FT, Li YM, Kounnas MZ, Thinakaran G (2007) Dual roles of the transmembrane protein p23/TMP21 in the modulation of amyloid precursor protein metabolism. Mol Neurodegener 2: 4

Walker A, Su H, Conti MA, Harb N, Adelstein RS, Sato N (2010) Non-muscle myosin II regulates survival threshold of pluripotent stem cells. Nat Commun 1: 71

Wang Y, Zhou Z, Walsh CT, McMahon AP (2009) Selective translocation of intracellular Smoothened to the primary cilium in response to Hedgehog pathway modulation. Proc Natl Acad Sci U S A 106: 2623–8

Washington Smoak I, Byrd NA, Abu-Issa R, Goddeeris MM, Anderson R, Morris J, Yamamura K, Klingensmith J, Meyers EN (2005) Sonic hedgehog is required for cardiac outflow tract and neural crest cell development. Dev Biol 283: 357–72

Wu X, Walker J, Zhang J, Ding S, Schultz PG (2004) Purmorphamine induces osteogenesis by activation of the hedgehog signaling pathway. Chem Biol 11: 1229–38

Wurth L, Papasaikas P, Olmeda D, Bley N, Calvo GT, Guerrero S, Cerezo-Wallis D, Martinez-Useros J, Garcia-Fernandez M, Huttelmaier S, Soengas MS, Gebauer F (2016) UNR/CSDE1 Drives a Post-transcriptional Program to Promote Melanoma Invasion and Metastasis. Cancer Cell 30: 694–707

Wutz A, Jaenisch R (2000) A shift from reversible to irreversible X inactivation is triggered during ES cell differentiation. Mol Cell 5: 695–705

Xiao X, Tang JJ, Peng C, Wang Y, Fu L, Qiu ZP, Xiong Y, Yang LF, Cui HW, He XL, Yin L, Qi W, Wong CC, Zhao Y, Li BL, Qiu WW, Song BL (2017) Cholesterol Modification of Smoothened Is Required for Hedgehog Signaling. Mol Cell 66: 154–162 e10

Xiong H, Zhao W, Wang J, Seifer BJ, Ye C, Chen Y, Jia Y, Chen C, Shen J, Wang L, Sui X, Zhou J (2017) Oncogenic mechanisms of Lin28 in breast cancer: new functions and therapeutic opportunities. Oncotarget 8: 25721–25735

Yam PT, Langlois SD, Morin S, Charron F (2009) Sonic hedgehog guides axons through a noncanonical, Src-family-kinase-dependent signaling pathway. Neuron 62: 349–62

Yilmaz A, Peretz M, Aharony A, Sagi I, Benvenisty N (2018) Defining essential genes for human pluripotent stem cells by CRISPR-Cas9 screening in haploid cells. Nat Cell Biol 20: 610–619

Ying QL, Wray J, Nichols J, Batlle-Morera L, Doble B, Woodgett J, Cohen P, Smith A (2008) The ground state of embryonic stem cell self-renewal. Nature 453: 519–23

Yuan X, Cao J, He X, Serra R, Qu J, Cao X, Yang S (2016) Ciliary IFT80 balances canonical versus non-canonical hedgehog signalling for osteoblast differentiation. Nat Commun 7: 11024

Zhang XM, Ramalho-Santos M, McMahon AP (2001) Smoothened mutants reveal redundant roles for Shh and Ihh signaling including regulation of L/R symmetry by the mouse node. Cell 106: 781–92

